# A role for APP in the development of astrocyte morphological complexity

**DOI:** 10.1101/2023.09.09.556967

**Authors:** Margaux Saint-Martin, Yukiko Goda

## Abstract

The amyloid precursor protein (APP), whose proteolytic cleavage gives rise to amyloid-*β* peptide, has been extensively studied for its role in Alzheimer’s disease, but its physiological function remains less understood. In neurons, APP and its two homologs, the amyloid precursor-like protein 1 (APLP1) and 2 (APLP2), are present in the synaptic compartment and promote synaptogenesis. Over recent years, astrocytes, an abundant glial cell in the brain, have attracted much attention for their role in regulating synapse formation and function. Although APP is also found in astrocytes, its role in these cells remains largely unexplored. Here we have sought to investigate the expression and function of APP in rodent astrocytes *in vitro* and *in vivo*. We find that APP along with its family members, APLP1 and APLP2, are expressed in astrocytes *in vitro* and *in vivo*. In primary hippocampal cultures, shRNA-mediated knockdown of astrocytic APP, APLP1 or APLP2 compromises astrocyte morphological elaboration to varying degrees. We have then focused on APP to characterize its role in astrocyte development in the intact brain. We show that astrocytic APP shapes the morphological complexity of astrocyte processes *in vivo* and may also modulate postsynaptic assembly. Our results highlight a role of astrocytic APP and possibly of APLPs that could potentially impact neuronal functions.

## Introduction

The amyloid precursor protein (APP), whose cleavage product is implicated in the pathogenesis of Alzheimer’s disease, has been the target of extensive research (Selkoe and Hardy, 2016), but its physiological function has been less well studied. APP gene family includes the amyloid precursor like protein 1 (APLP1) and 2 (APLP2) that share common structural and functional features with APP. APP and APLPs are cell adhesion proteins with a transmembrane domain, and their processing by secretases generates various intracellular and extracellular fragments (Müller et al., 2017). All three proteins have been found to play partially redundant functions in the nervous system, and accordingly single KO mice deficient in genes encoding any one of these proteins are viable. Notably, however, APP/APLP2 and APLP1/APLP2 double KO mice die shortly after birth (Heber et al., 2000), whereas APP/APLP1 double KO mice are viable. These observations suggest that APP and APLP1 might have non-overlapping roles in the nervous system that are compensated by the presence of APLP2.

Aside from its involvement in the ageing brain, APP has been shown to play a role in developmental processes including neuronal migration, neurite growth and synapse formation (Perez et al., 1997; Priller et al., 2006; Young-Pearse et al., 2007, 2008; Southam et al., 2019). The expression of APP, APLP1 and APLP2 proteins reaches a peak between 1 and 2 weeks after birth, consistent with a major role of these proteins during development (Trapp and Hauer, 1994; Schilling et al., 2017). Of note, APP family members interact with each other in both “*cis*” (on the same membrane surface) and “*trans*” (intercellularly across two membrane surfaces) in homo- and heterophilic fashion, and *trans* interactions in particular are thought to promote cell adhesion (Soba et al., 2005; Stahl et al., 2014; Schilling et al., 2017). That APP family members possess synaptic adhesion molecule (SAM)-like functions is further supported by studies in which APP family members are shown to induce presynaptic differentiation with APP and APLP1 displaying high efficacy (Wang et al., 2009; Stahl et al., 2014; Schilling et al., 2017). This synaptogenic activity is suggested to rely on intercellular *trans* interactions involving presynaptic APP, APLP1 or APLP2(Wang et al., 2009; Schilling et al., 2017).

Recent reports have demonstrated the expression of diverse SAMs in astrocytes such as ephrins or neuroligins (Hillen et al., 2018; Tan and Eroglu, 2021; Saint-Martin and Goda, 2022). Astrocytic SAMs facilitate the interaction of astrocyte processes with synapses and promote astrocyte and synapse development. Interestingly, APP has also been found in astrocytes (Haass, Hung and Selkoe, 1991; LeBlanc et al., 1991) where it has been implicated in lipoprotein metabolism, Ca^2+^ homeostasis and mitochondrial function (Linde et al., 2011; Fong et al., 2018; Montagna et al., 2019). However, whether astrocytic APP plays a role in astrocyte and synapse development remains unknown. In this study we evaluate the expression of APP along with its family members APLP1 and APLP2 in astrocytes, and its function in astrocyte and synapse development *in vitro* and *in vivo*.

## Material and Methods

### Cloning

PCR amplifications for cloning were performed with the CloneAmp HiFi PCR Premix (Clontech) and plasmid fusion/reassembly were performed with the NEBuilder HiFi DNA Assembly Master Mix (NEB). To create the empty shRNA plasmid (shE), the shRNA sequence from the pU6-shAPP plasmid was deleted by using PCR amplification of the whole vector outside the shAPP sequence followed by reassembly. For the scramble shRNA (GACCGATCCACATTGTCGTAA), a scramble sequence of the shAPP sequence was generated with Invivogen scramble siRNA Wizard software. This sequence should not match with any rat or mouse mRNA. Complementary DNA oligomers for the scramble shRNA sequences with ends overlapping with the receiving vector (Thermofisher Scientific) were annealed to generate double stranded oligomers. These oligomers were then inserted into the pENTR-U6 vector by fusion.

To create the AAV-pGFAP104-hAPP751 Ct-HA plasmid, the EGFP insert from the AAV-pGFAP104-EGFP plasmid was replaced by the hAPP695 Ct-HA sequence from the pCMV-hAPP695 Ct-HA plasmid kindly provided by Drs. Simone Eggert and Edward Koo. The hAPP695 sequence was then replaced with the hAPP751 sequence from the pGEM-hAPP751 plasmid (also kindly provided by Drs. Simone Eggert and Edward Koo). The pU6-shRNA plasmids against APP, APLP1 and APLP2 from Young-Pearse (2007 and 2008) were obtained from Addgene. The target sequences were as follows: shAPP (GCACTAACTTGCACGACTATG), shAPLP1 (CGTAGGATGCGCCAGATTAA), shAPLP2 (GATATGGACCAGTTTACC). AAV-pGFAP-EGFP-shRNA(miR30) plasmids (shEmpty, shSbl and shAPP) were designed by VectorBuilder with the same target sequences as the pU6-shEmpty, pU6-shSbl and pU6-shAPP plasmids. The miR30 sequences flanking the shRNA sequence allow the expression of shRNAs under a GFAP polII promoter instead of a polIII promoter such as U6.

### HEK cultures and transfection - Antibody and shRNA validation

Human embryonic kidney (HEK) 293FT cells (ThermoFisher Scientific) were grown at 37°C, 5% CO_2_ in DMEM containing 10% foetal bovine serum (FBS), 1% Penicillin/Streptomycin (P/S) and regularly passaged every three days for maintenance. For antibody and shRNA validation 80 000 cells were plated on 12mm glass coverslips coated with poly-D-lysine (PDL) placed in 24 well plates or 300,000 cells were plated in 3.5cm petri dishes. Cells were then transfected with lipofectamine 2000 (Invitrogen) following the manufacturer’s instructions. To validate antibodies against APP, APLP1 and APLP2, HEK cells were transfected with pCMV-hAPP695 Ct-HA, pCMV-hAPLP1 Ct-HA or pCMV-hAPLP2 Ct-HA. After 48 h cells were fixed with 4% PFA for immunofluorescence experiments or lysed in NaCl 150 mM, HEPES 50 mM pH 7.5, Triton X-100 1% with protease and phosphatase inhibitors for western blot analyses. Labelling with antibodies against the HA-tag and against each protein confirmed the ability of APP, APLP1 or APLP2 antibodies to specifically bind to their target (Fig. S1). To verify shRNAs’ efficacy, HEK cells were transfected with plasmids containing the target protein mAPP, hAPLP1 or mAPLP2 along with their respective shRNA (shAPP, shAPLP1 or shAPLP2) or control empty (shE) or scramble (shSbl) shRNAs and GFP. Two days after transfection, HEK cells were either fixed for immunofluorescence labelling or lysed for western blot analyses. Knockdown was confirmed by comparing APP, APLP1 or APLP2 levels in shAPP, shAPLP1 or shAPLP2 conditions respectively to the control shE and shSbl conditions (Fig. S2A, B). Plasmids expressing APP, APLP1 or APLP2 were all kindly provided by Drs. Simone Eggert and Edward Koo.

### Primary hippocampal astrocyte and neuronal cultures and co-cultures

For all primary hippocampal cultures, hippocampi from P0-P1 rats were dissected and digested in 1.5 mM CaCl_2_, 0.15 mg/ml L-Cysteine, 0.5 mM EDTA, 1 µg/ml DNAse I, 20 units/ml of papain for 20 min at 37°C. Hippocampi were then washed and triturated in astrocyte culture medium (BME, 10% FBS, 20 mM glucose, 1 mM Na-pyruvate, 1X P/S). All cells were kept at 37°C in 10% CO_2_.

### Primary hippocampal neuronal cultures

For neuronal cultures, 60 000 cells were plated onto 12 mm glass coverslips coated with 10 µg/ml poly-D-lysine (PDL) and 4 µg/ml laminin in 24 well plates with astrocyte culture medium. The medium was then changed to neuronal culture medium (Neurobasal, 1X glutamax, 20 mM glucose, 1X B27, 1 µg/ml MITO^TM^ serum extender, 1X P/S) after 5-6 h incubation at 37°C, 10% CO_2_. On day *in-vitro* (DIV) 4, 4 µM AraC was added to the neuronal culture medium to stop the proliferation of astrocytes. Cells were fixed at DIV 18 with 4% PFA, 4% sucrose in PBS at room temperature (RT) for 10 min or with ice cold methanol for 15 min on ice.

### Astrocyte culture

For astrocyte cultures, cells in astrocyte culture medium were either plated onto 12mm glass coverslips coated with 4 µg/ml PDL + 1 mg/ml collagen at 15,000 cells per well in a 24 well plate, or on a 25 mm^2^ flask with 1 million cells. Astrocytes in 24 well plates were kept at 37°C, 10% CO_2_ until cell fixation (4% PFA, 4 % sucrose in PBS) at DIV 18 and used for immunostaining. Astrocytes in flasks were passaged at DIV11 with trypsin-EDTA 0.05% to 3.5 cm petri dishes and transfected at DIV13 with Lipofectamine-2000. At DIV15, astrocytes were either lysed with NaCl 150 mM, HEPES 50 mM pH 7.5, Triton X-100 1% with protease and phosphatase inhibitors for western blot or detached with trypsin-EDTA 0.05%, centrifuged, and resuspended in neuronal culture medium to be plated on top of neuronal cultures at 15,000 astrocytes per well. At this stage, cell counting of both total number of cells and fluorescent cells allowed us to estimate the transfection efficacy at approximatively 10-20% (data not shown). Astrocyte-neuron co-cultures were then fixed at DIV18 with 4% PFA, 4% sucrose in PBS for 10 min at RT and used for immunofluorescence labelling.

### HEK-Astrocyte co-cultures - substrate experiment

For substrate experiments, HEK cells were grown in 3.5 cm petri dishes and transfected with lipofectamine 2000 to express HA tagged Neurexin-1β (NRXN-1β-HA), neuroligin 1 (NL1-HA), APP (APP-HA), APLP1 (APLP1-HA), APLP2 (APLP2-HA) or mGFP. Two days after transfection, HEK cells were lifted and plated on 12 mm glass coverslips coated with 10 µg/ml PDL in 24 well plates at 250,000 cells per well. The next day, non-transfected astrocytes were plated on top of HEK cells at 5000 cells per well and left to grow for 24 h before fixing the cells with 4% PFA in PBS. Cells were then immunolabeled with anti-HA antibodies to identify transfected HEK cells and with anti-GFAP antibodies to visualize astrocyte branches.

### Western Blot

For western blot, cells grown in 3.5 cm petri dishes (HEK cells, primary neuronal or astrocyte culture or astrocytes transfected with shRNAs) were lysed in NaCl 150 mM, HEPES 50 mM pH 7.5, Triton X-100 1% with protease and phosphatase inhibitors for 10 min on ice. Cell lysates were then centrifuged at 12,000 x g for 15 min at 4°C. Pellet containing debris was then discarded and lysate supernatant was diluted in 1X Laemmli buffer, 50 mM DTT and heated to 95°C for 5 min. Proteins were then separated on 7.5% polyacrylamide gels and transferred to a PVDF membrane. Antibodies against APP, APLP1 or APLP2 were used to evaluate the protein respective expression level and antibodies against β-actin were used as a loading control. Primary antibodies were labelled with Horseradish Peroxidase (HRP) secondary antibodies and revealed with Amersham ECL prime (Cytiva) on a Vilber Fusion X machine.

### Immunofluorescence on primary cell cultures

For cell culture immunolabelling, cells were permeabilized and blocked for 30 min in phosphate-buffered saline (PBS), 0.3% Triton X-100, 1% bovine serum albumin (BSA), 10% normal goat serum (NGS) before incubating with primary antibodies in PBS, 0.3% Triton X-100, 1% BSA, 4% NGS for 2 hours at RT. Cells were then washed in PBS 3 times and incubated with secondary antibodies in PBS, 0.3% Triton X-100, 1% BSA, 4% NGS for 1 hour at RT. After 3 washes in PBS, coverslips were mounted on glass slides with Vectashield (Vector laboratories), 1/1000 DAPI and sealed with nail polish. Images were acquired on an inverted confocal laser scanning microscope Olympus FV3000 at the RIKEN CBS-EVIDENT Open Collaboration Center (BOCC) and RIKEN Center for Brain Science, Research Resources Division (RRD). Reference and dilutions of antibodies used are in the antibody table.

### *In vivo* experiments

#### Neonatal sinus injection method (n-SIM)

For brain AAV injections, n-SIM method described by Hamodi et al. (2020) was used. Recombinant AAV vectors were prepared according to standard procedures (Grieger et al., 2006). AAV9-pGFAP-EGFP-shE, AAV9-pGFAP-EGFP-shSbl or AAV9-pGFAP-EGFP-shAPP were diluted in PBS to a concentration of 4x10^12^ vg/ml. P0-P1 ICR mice were anesthetized on ice for 5-10 min. A 10 µl NanoFil syringe (WPI, NANOFIL) with 35G bevelled needles (WPI, NF35BV-2) was filled. Pups were placed on a homemade cold plate on a stereotaxic table (Muromachi, 51500) and small cuts were performed above each sinus. The syringe was then lowered into the sinus to penetrate and positioned about 300-500 µm under the surface of the skull. For each sinus, 2 µl of AAV was injected over a period of 100 s followed by a 5 s pause before removal of the syringe. In total, each mouse received 4µl of AAV. The skin was then unfolded back in place and the surface was cleaned with a Q-tip before placing the pups on a warm pad to recover.

#### Paraformaldehyde fixation

After three weeks, mice were anesthetized with an intraperitoneal injection of 0.3 mg/ml medetomidine, 4 mg/kg midazolam, 5 mg/kg butorphanol. Mice were then perfused with PBS followed by 4% PFA in PBS. Fixed brains were retrieved and placed in 4% PFA at 4°C overnight. Brains were then incubated in 20% sucrose in PBS at 4°C overnight followed by incubation in 30% sucrose in PBS at 4°C overnight before freezing them in Tissue-Tek OCT.

#### Brain slices and IHC

Floating sections were prepared on a cryostat to a 20 µm or 100 µm thickness and kept at -20°C in antifreeze medium (20% Glycerol, 30% Ethylene glycol, 50% PBS). For immunohistochemistry, 20 µm thick brain slices were used. Briefly, slices were washed in PBS before incubating in blocking solution (PBS, 0.3% Triton X-100, 1% BSA, 10% NGS) for 1 h at RT. Brain slices were then incubated with primary antibodies diluted in PBS, 0.3% Triton X-100, 1% BSA, 4% NGS at 4°C overnight. Slices were then washed in PBS, 0.3% Triton X-100 and incubated with secondary antibodies and DAPI diluted in PBS, 0.3% Triton X-100, 1% BSA, 4% NGS for 1-2 h at RT. Slices were then washed in PBS and mounted on glass slides with vectashield. Images were acquired on an inverted confocal laser scanning microscope Olympus FV3000 at the RIKEN CBS-EVIDENT Open Collaboration Center (BOCC) and RIKEN Center for Brain Science, Research Resources Division (RRD). Reference and dilutions of used antibodies are in the antibody table. Secondary antibody-only labellings were performed for each secondary antibody used, and did not show any non-specific staining.

### Analysis

#### Sholl analysis

The complexity of astrocyte morphology *in vitro* was analysed using the ImageJ Sholl analysis plugin on confocal (Olymus FV3000) images taken with a 20x air or 40x silicon immersion objective. The cell soma center was assessed by using the DAPI staining and circles were traced every 1 µm from the center of the cell soma to the furthest process tip. The number of intersections between astrocyte processes and each circle was then counted. Values were summed by 10 µm brackets for representation and statistical analysis.

#### Synapse number analysis *in vitro*

Synapse number *in vitro* was analysed using the ImageJ plugin Puncta Analyzer (written by Bary Wark) (Ippolito and Eroglu, 2010) on confocal images taken with a 60x silicon immersion objective. Confocal z-stack images were taken with a 0.5 µm step and analysis was performed on maximum intensity projections. The number of Homer1, VGLUT1, and co-localized puncta (synapse) inside the astrocyte area was measured in each condition. To isolate one astrocyte area other astrocyte processes in the image were erased and binary images of the target astrocyte area were obtained.

#### *In vivo* APP expression analysis

Knockdown of APP expression in astrocytes *in vivo* was assessed using the 3D analysis software, IMARIS. Briefly, astrocytes were 3D reconstructed using the IMARIS surface tool based on their GFP expression. All APP signal outside the GFP-labelled astrocyte surface was then deleted and the remaining APP signal was reconstructed. The volume of both the astrocyte and the APP signal inside were measured. The efficacy of APP KD was then evaluated by comparing the volume of APP signal per astrocyte volume between the shSbl and shAPP conditions. Values were normalized to the control shSbl condition for each experiment. This analysis was performed at least on 5 astrocytes from every section for each mouse (6 mice per condition).

#### Neuropil infiltration volume (NIV) analysis

Confocal z-stack images were taken using a 60x silicon immersion objective with a 2x optical zoom and a 0.5 µm step from 20 or 100 µm brain sections. Images were then analysed on IMARIS for 3D visualisation. The NIV is a measure of the volume of astrocyte processes that can vary depending on both astrocyte process number and size. For each astrocyte, 3 regions of 4 µm x 4 µm x 4 µm (64 µm^3^) were randomly selected that were away from any major branches or cell soma (determined by DAPI staining). Astrocyte processes in ROI were reconstructed using the IMARIS surface tool and their volume was measured.

#### Astrocyte volume analysis

Confocal z-stack images were taken using a 60x silicon immersion objective with a 2x optical zoom and a 0.5 µm step from 100 µm brain sections. Only astrocyte with their entire cell body within the z-stack were selected for analysis. Astrocytes were reconstructed in 3D using the IMARIS surface tool and total astrocyte volume was measured.

#### Astrocyte territory analysis

Astrocyte territory analysis was performed on the same images as the astrocyte volume analysis, using ImageJ on maximum intensity projections. Astrocyte territory was traced manually and the area was obtained using the measure tool.

#### Synapse number analysis *in vivo*

Synapse number *in vivo* was analysed using the ImageJ plugin Puncta Analyzer (written by Bary Wark) on confocal images taken with a 60x silicon immersion objective with a 2x optical zoom and a 0.5 µm step. Analysis was performed on maximum intensity projections from two optical sections for each astrocyte (5 astrocytes/slice, 2 slices/mouse, 3 mice/condition). The number of postsynaptic (homer1 or gephyrin), presynaptic (VGLUT1 or VGAT), and co-localized puncta (synapses) was measured in 3 randomly selected regions of 7 µm x 7 µm (49 µm^2^) inside and outside the GFP positive astrocyte area. A ratio of puncta inside target astrocyte area over outside target astrocyte area was then calculated for each condition.

### Statistical tests

All statistical analyses were performed on GraphPad Prism 9. Parametric tests were used when data was normal, otherwise non-parametric test were used. Sample size and statistical test used are indicated in the figure legend for each experiment; p-values for significant differences are detailed in the text. Significance is noted as follows: * p ≤ 0.05; ** p ≤ 0.01; *** p ≤ 0.001; **** p ≤ 0.0001.

### Antibody list (Supplementary Table 1)

| Antibody | Species | Company | Reference | Dilution IF | Dilution WB |
| --- | --- | --- | --- | --- | --- |
| **APP** | Rabbit | Abcam | Ab32136 (Y188) | 1/200-1/300 | 1/800-1/1000 |
| **APLP1** | Rabbit | ThermoFisher | PA5-78795 | 1/200-1/300 | 1/800-1/1000 |
| **APLP2** | Rabbit | Abcam | Ab140624 (EPR5938) | 1/200-1/300 | 1/500 |
| **GFP** | Rabbit | Millipore | AB3080P | 1/500-1/1000 | 1/1000-1/2000 |
| **GFP** | Mouse | abcam | ab1218 (9F9.F9) | 1/500-1/1000 | 1/1000-1/2000 |
| **GFAP** | Rabbit | abcam | ab7260 | 1/500-1/1000 |  |
| **GFAP** | Mouse | Synaptic System | 173 011 (134B1) | 1/500-1/5000 |  |
| **HA** | Mouse | Sigma | H3663 | 1/500 | 1/1000 |
| **Homer1** | Rabbit | Synaptic System | 160 003 | 1/300 |  |
| **VGLUT1** | Guinea Pig | Synaptic System | 135 304 | 1/500 |  |
| **Gephyrin** | Mouse | Synaptic System | 147 011 (mAb7a) | 1/300 |  |
| **VGAT** | Guinea Pig | Synaptic System | 131 004 | 1/500 |  |
| **Synaptophysin** | Guinea Pig | Synaptic System | 101 004 | 1/500-1/1000 |  |
| **β-actin** | Mouse | Sigma | A1978 (AC-15) |  | 1/10000 |
| **Alexa Fluor 488 Goat anti-Mouse** | goat | Invitrogen | A11029 | 1/500-1/1000 |  |
| **Alexa Fluor 488 Goat anti-Rabbit** | goat | Invitrogen | A11034 | 1/500-1/1000 |  |
| **Alexa Fluor 594 Goat anti-Mouse** | goat | Invitrogen | A11032 | 1/500-1/1000 |  |
| **Alexa Fluor 594 Goat Anti-Rabbit** | goat | Invitrogen | A11012 | 1/500-1/1000 |  |
| **Alexa Fluor 633 Goat anti-Mouse** | goat | Invitrogen | A21052 | 1/500-1/1000 |  |
| **Alexa Fluor 647 Goat anti-Guinea Pig** | goat | Invitrogen | A21450 | 1/500-1/1000 |  |
| **DAPI, dilactate** |  | SIGMA | D9564 | 1/500-1/1000 |  |
| **Anti-Mouse Peroxidase antibody** | rabbit | Sigma | A9044 |  | 1/6000 |
| **Anti-Rabbit Peroxidase antibody** | goat | Sigma | A0545 |  | 1/6000 |

### AAV and plasmid list (Supplementary Table 2)

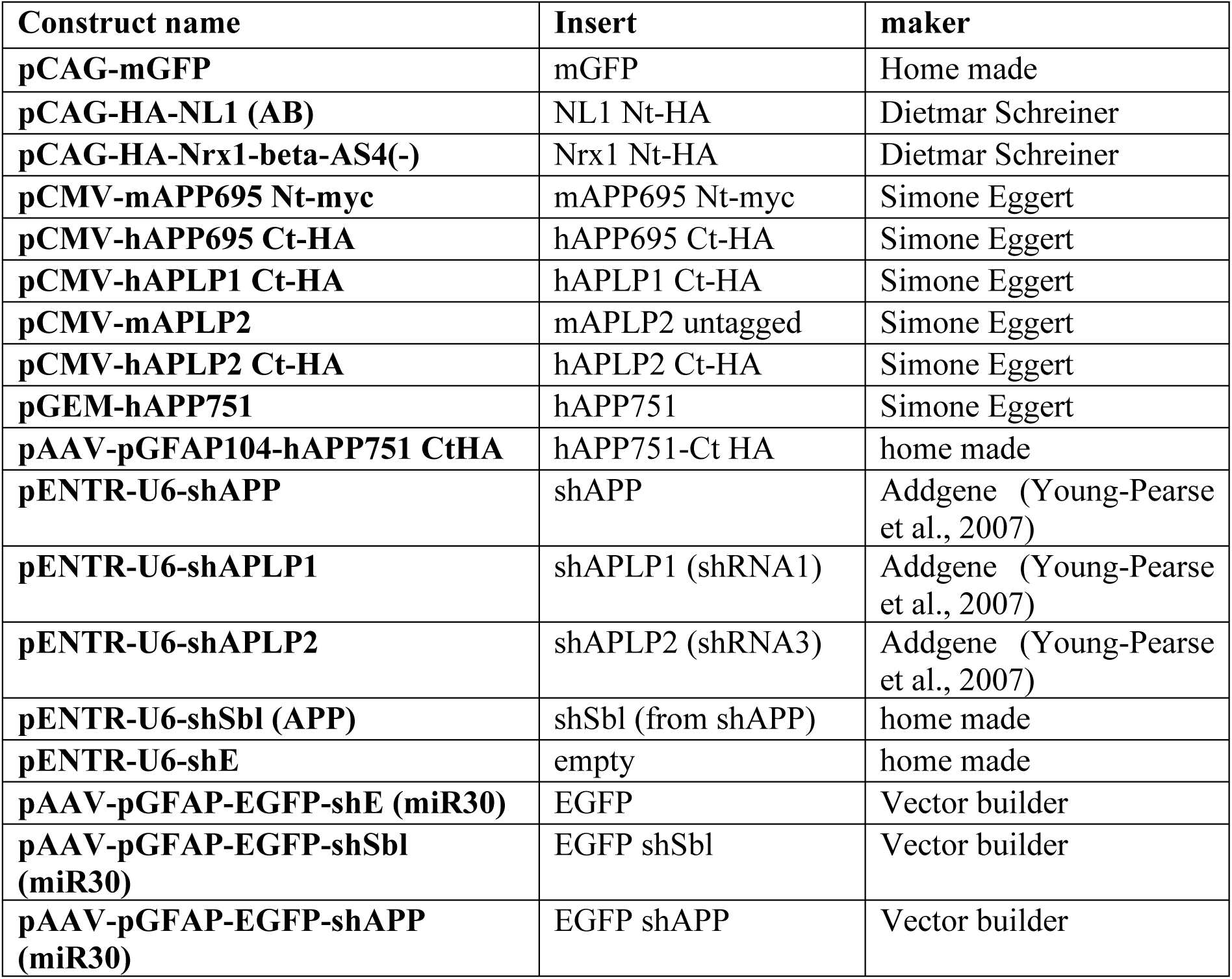

## Results

### APP and APLPs are expressed in neurons and astrocytes

Contrary to the well accepted expression of APP and its family members APLP1 and APLP2 in neurons (LeBlanc et al., 1991; Lorent et al., 1995; Schilling et al., 2017), their expression in astrocytes has remained unclear with conflicting evidence (Haass, Hung and Selkoe, 1991; LeBlanc et al., 1991; Guo et al., 2012; Del Turco et al., 2016). While APP was present in astrocytes in culture (Haass, Hung and Selkoe, 1991; LeBlanc et al., 1991), APP could not be detected in astrocytes *in vivo* (Guo et al., 2012; Del Turco et al., 2016). A recent patch-Seq transcriptome analysis of mice hippocampal astrocytes in our team revealed the presence of App, Aplp1 and Aplp2 mRNA in astrocytes in area CA1 (data not shown). Upon searching the publicly available rodent brain transcriptomic (Fig. S1A) and translatomic (Fig. S1B) datasets, App, Aplp1 and Aplp2 mRNA were repeatedly found in astrocytes in the cortex (Cahoy et al., 2008; Zhang et al., 2014; Srinivasan et al., 2016; Sakers et al., 2017) and in the hippocampus (Clarke et al., 2018a; Mazaré et al., 2020). Furthermore, App, Aplp1 and Aplp2 mRNA were also detected in perisynaptic astrocyte processes (PAPs) in translating ribosome affinity purification (TRAP) experiments, suggesting that they could be actively translated locally in PAPs (Sakers et al., 2017; Mazaré et al., 2020). We summarized the transcriptomic and translatomic datasets from various publications for App, Aplp1, Aplp2 in Supplemental Figure 1, in which mRNA levels for neuroligin 2 (Nlgn2), another synaptic cell adhesion molecule expressed in astrocytes, were included also for reference. Overall, App, Aplp1 and Aplp2 mRNA were expressed in astrocytes at a level similar or lower compared to their respective expression in neurons (Cahoy et al., 2008; Zhang et al., 2014). Notably, in most databases, levels of App, Aplp1 or Aplp2 mRNA in astrocytes were higher than the levels of Nlgn2 mRNA in astrocytes. Interestingly, Clarke et al. (2018) found in the hippocampus that astrocytic Aplp1 and Aplp2 mRNA expression strongly increased across the age of animal whereas astrocytic App mRNA expression remained constant (Fig. S1B); in the cortex, all three gene expression in astrocytes increased with age (Clarke et al., 2018a).

Given the data supporting the mRNA expression of App, Aplp1 and Aplp2 in astrocytes *in vitro* and *in vivo*, we began our study by confirming the protein expression of APP, APLP1 and APLP2 in astrocytes relative to their expression in neurons *in vitro*. We used two types of DIV18 primary cultures: neuron-astrocyte co-cultures and astrocyte enriched cultures. Cultures were either fixed with cold methanol and processed for immunolabelling or lysed for western blot analyses. Antibodies against APP, APLP1 and APLP2 were validated on HEK cells overexpressing HA-tagged APP, APLP1 or APLP2 protein (Fig. S2A and 2B). GFAP labelling was used to identify astrocytes, and synaptophysin was used as a neuronal synapse marker (Fig. 1A, 1B). APP, APLP1 and APLP2 immunofluorescence signals were detected not only in neurons (Fig. 1A) but also in astrocytes (Fig. 1A, B). In neuron-astrocyte co-cultures, APP, APLP1 and APLP2 immunofluorescence signals were overall weaker in astrocytes compared to signals associated with neurons (Fig. 1A). Consistently, in western blot experiments, APP, APLP1 and APLP2 levels were lower in astrocyte-enriched cultures compared to their levels in neuron-astrocyte co-cultures (Fig. 1C).

**Figure 1:**
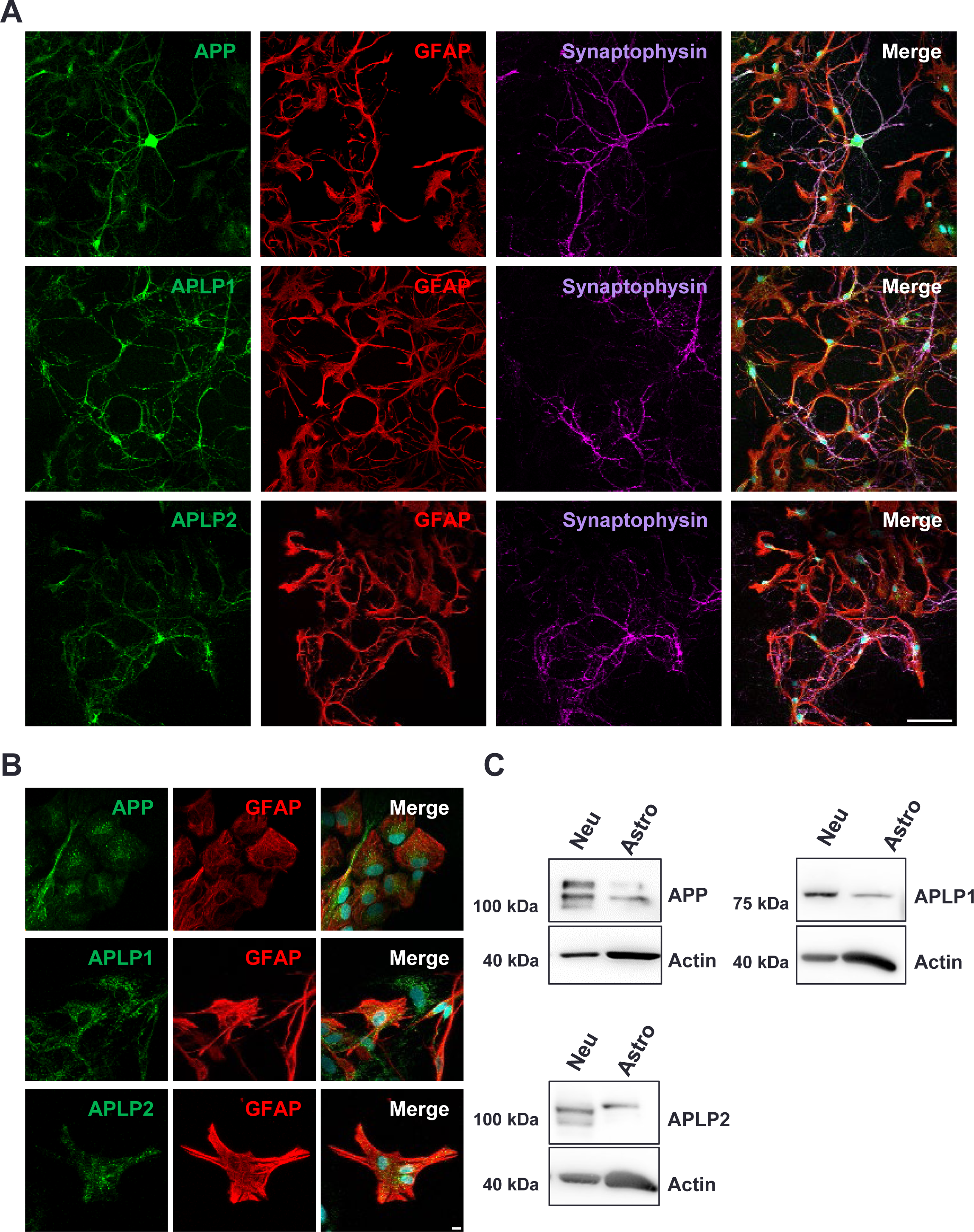
APP, APLP1 and APLP2 expression in neuron-astrocyte co-cultures and in astrocyte cultures. **A)** DIV18 neuron-astrocytes co-cultures were stained with APP, APLP1 or APLP2 antibodies. GFAP antibody was used to label astrocytes and synaptophysin antibody was used to label synapses. Images were taken on a confocal with a 10x objective. Scale bar 100 µm. **B)** DIV 18 astrocyte cultures were stained with APP, APLP1 or APLP2 antibodies along with GFAP antibodies. Images were taken on a confocal with a 20x objective. Scale bar 10 µm. **C)** Western Blot analysis of DIV18 neuron-astrocyte co-cultures and astrocyte cultures. Actin antibody was used as a loading control.

To examine APP, APLP1 and APLP2 protein expression in astrocytes *in vivo*, mice were injected with AAV9 to drive the expression of EGFP under the GFAP promoter (AAV-pGFAP-EGFP-shE(miR30) or shSbl(miR30)) (Fig 2A, B and S3A, B). The GFP signal allowed the visualization of the entire astrocyte including PAPs and small astrocyte leaflets while synaptophysin was used to locate the presynaptic compartment of neurons (Fig. 2C and S3C). APP, APLP1 and APLP2 labelling could be found inside GFP positive regions, suggesting their presence in hippocampal and cortical astrocyte processes. Of note, because of the close proximity of synapses to astrocyte processes and the previously demonstrated synaptic expression of APP, APLP1 and APLP2 in neurons, we could not exclude the possibility that at least part of the overlapping APP, APLP1 and APLP2 signals with astrocyte processes came from a neighbouring synapse or a neuron. Overall, in line with previous literature, our data are consistent with the expression of APP protein in astrocytes *in vitro* and *in vivo*. Moreover, APLP1 and APLP2 proteins are also expressed in astrocytes *in vitro* and *in vivo*. However, the expression of all three proteins in astrocytes are at levels lower than that found in neurons.

**Figure 2:**
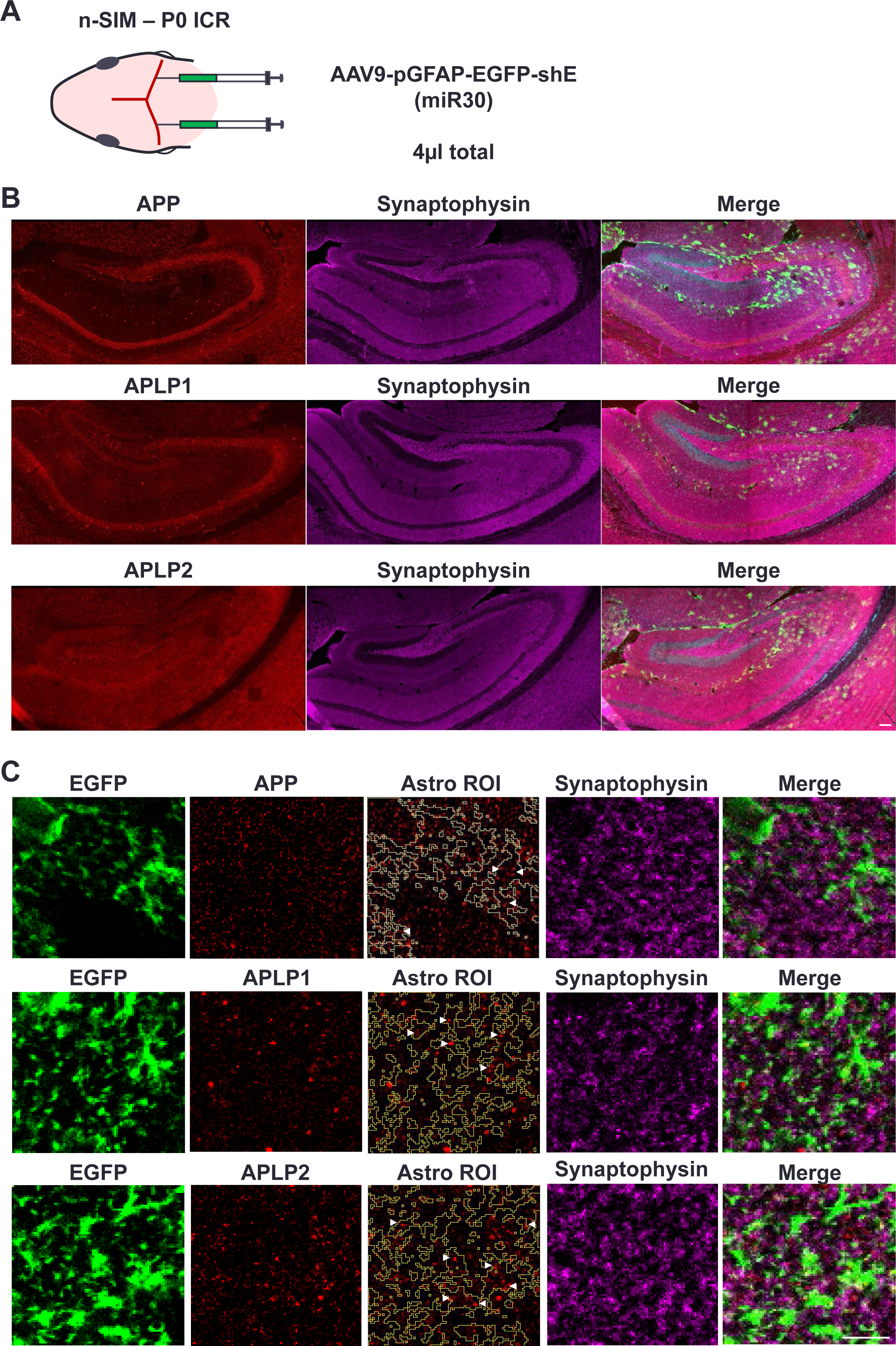
APP, APLP1 and APLP2 expression *in vivo*. **A)** Neonatal sinus injection method. AAV with EGFP and an empty shRNA under the GFAP promoter for astrocyte specific expression was injected in newborn ICR mice at P0. **B)** Hippocampus from a P21 mouse injected with AAV9-pGFAP-EGFP-shE(miR30) and labelled with antibodies against synaptophysin and APP, APLP1 or APLP2. Mosaic images were taken on a confocal microscope with a 10x objective. Sparse astrocytic EGFP expression can be observed. **C)** Localisation of APP, APLP1 and APLP2 in astrocyte processes. Hippocampal astrocytes (CA3 SR) were imaged on a confocal microscope with a 60x silicon immersion objective and a 2x optical zoom, only one optical plane is presented here. The EGFP expressed in astrocytes allows the visualization of small astrocyte processes and synaptophysin allows the localisation of synapses. Astrocyte processes are in close contact with synaptophysin puncta. APP, APLP1 and APLP2 puncta are inside (arrowheads) and outside the EGFP positive astrocyte ROI (Astro ROI). Scale bar 5µm

### Astrocytic APP and APLPs impact astrocyte morphology *in vitro*

APP was shown to play a role in neuronal development especially on neurite outgrowth (Perez et al., 1997; Young-Pearse et al., 2008; Weyer et al., 2014). Given the presence of APP in astrocytes, we thus wondered if APP and also its related family members APLP1 and APLP2 could play a role in astrocyte development. To test this hypothesis, we knocked down APP, APLP1 or APLP2 in primary rat hippocampal astrocytes using shRNAs (Young-Pearse et al., 2007, 2008). As controls, an empty shRNA (shEmpty) and a scramble shRNA (shSbl) generated from the shAPP sequence were used. The efficacy of shRNAs was validated in HEK cells exogenously overexpressing the APP, APLP1 or APLP2 protein (Fig. S4A, B) and in primary cultures of rat hippocampal astrocytes (Fig. S4C). To study the impact of APP, APLP1 or APLP2 knockdown (KD) on astrocyte morphological elaboration, shRNAs were co-transfected with a plasmid encoding membrane bound GFP (mGFP) to visualize astrocyte processes. Transfected astrocytes were then plated on top of neurons to let grow for additional 72 h – a condition that favoured the formation of morphologically complex astrocyte processes in cultures (Stogsdill et al., 2017) – before fixation (Fig 3A). Astrocyte morphological complexity (i.e. astrocyte branch complexity) was quantified by Sholl analysis that scored the number of processes intersecting with circumference of concentric circles drawn every micrometre starting from the cell soma to the furthest process (Fig 3B). Results were pooled by 10 µm bins and represented as number of intersections by a distance from the cell soma. We noted a significant decrease in the number of intersections in the shSbl expressing cells compared to the shEmpty condition. The expression of shRNA in astrocytes could potentially impact cell health or development since another shSbl sequence gave similar results (data not shown). Therefore, for statistical analysis the shSbl condition was used as the reference control. Relative to the shSbl condition, both APP KD and APLP1 KD significantly decreased astrocyte branch complexity (N = 6 cultures, repeated measures two-way ANOVA with Dunnett’s multiple comparison test at 30 and 40 µm distance; shSbl vs. shAPP, p < 0.0001; shSbl vs. shAPLP1, p < 0.0001) (Fig. 3B). Conversely, astrocyte complexity was significantly increased in APLP2 KD compared to the shSbl condition (N = 6 cultures, repeated measures two-way ANOVA with Dunnett’s multiple comparison test at 30 µm distance; shSbl vs. shAPLP2, p < 0.0017) (Fig. 3B, C). We also compared the effects of knocking down APP, APLP1 and APLP2 expression on the total process length and the area of astrocytes relative to controls. Interestingly, the total process length and the astrocyte area was decreased in the shAPLP1 but not in the shAPP condition (N = 6 cultures, One-way ANOVA with Dunnett’s multiple comparison test; shSbl vs. shAPLP1, total length and astrocyte area, p < 0.0001 and p = 0.0016, respectively) (Fig. 3C). Thus, although the number of processes was decreased in APP KD astrocytes, their total surface or total process length was not impacted. These results suggests that APP and APLPs modulate astrocyte process elaboration differentially *in vitro*.

**Figure 3:**
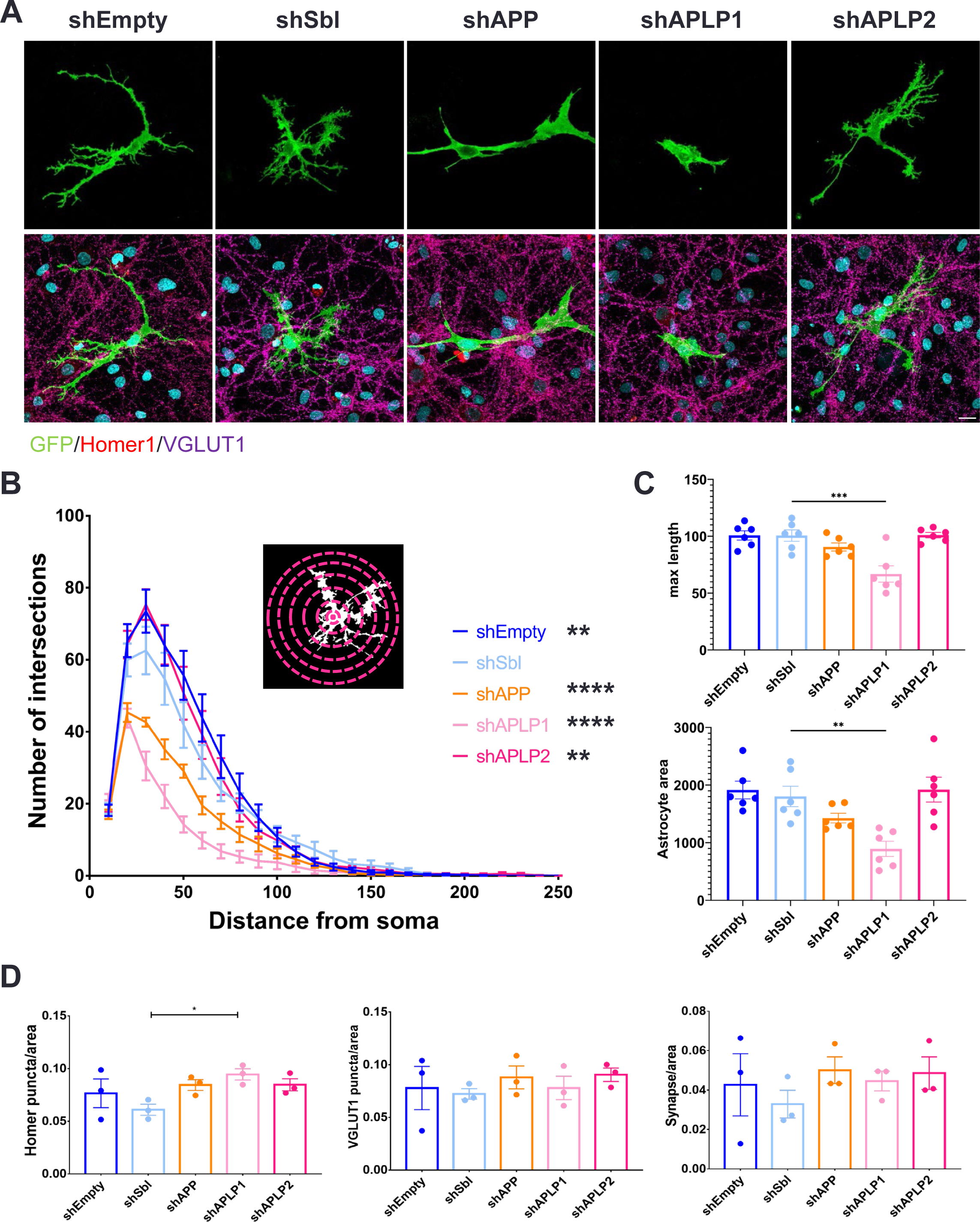
Impact of APP and APLPs on astrocyte morphology and synapses *in vitro*. **A)** DIV 18 astrocyte-neuron co-cultures. Astrocytes (green) were transfected with mGFP and shRNAs (shEmpty, shSbl, shAPP, shAPLP1 or shAPLP2) and grown on top of neurons at DIV15. Excitatory synapses were labelled with pre-synaptic VGLUT1 (purple) and post-synaptic Homer1 (red). Images were taken on a confocal microscope with a 60x silicon immersion objective. Scale bar 20µm. **B)** Analysis of astrocyte morphology complexity by Sholl analysis N=6 cultures (8-15 astrocytes per culture), repeated measures two-way ANOVA followed by Dunnett’s multiple comparison test. **C)** Quantification of astrocyte maximum process length and astrocyte area (N=6 cultures, 8-15 astrocytes per culture, One-way ANOVA followed by Dunnett’s multiple comparison test). **D)** Quantification of the number of Homer1, VGLUT1 and co-localized puncta per astrocyte area (N=3 cultures, 10 astrocytes per culture, One-way ANOVA followed by Dunnett’s multiple comparison test).

### Astrocytic APLP1 impacts excitatory post-synapse density *in vitro*

APP, APLP1 and APLP2 can interact homophilically and heterophilically both in *cis* and in *trans* to promote cell adhesion (Soba et al., 2005; Stahl et al., 2014; Schilling et al., 2017). Because of their expression in the presynaptic and the postsynaptic compartments (Schilling et al., 2017), astrocytic APP, APLP1 or APLP2 could potentially interact with their presynaptic or postsynaptic counterparts to form intercellular bridges at tripartite synapses. In addition, APP, APLP1 and APLP2 overexpressed in HEK cells could promote synapse formation in co-cultured neurons (Wang et al., 2009; Stahl et al., 2014; Schilling et al., 2017). We therefore investigated whether astrocytic APP, APLP1 and APLP2 could modulate synapse formation in our astrocyte-neuron co-cultures by knocking down the expression of APP, APLP1 or APLP2 specifically in astrocytes. Excitatory synapses were labelled with antibodies against VGLUT1 to mark presynaptic vesicles and Homer1 to mark the postsynaptic scaffold (Fig. 3A). We then counted puncta in each channel as well as co-localized presynaptic and postsynaptic puncta (i.e. synapses) inside an isolated astrocyte area (see Methods). Overall, no significant differences were observed between the conditions except for an increase in Homer1 puncta per astrocyte area upon APLP1 KD compared to the shSbl control (N = 3 cultures, 10 astrocytes per culture, One-way ANOVA with Dunnett’s multiple comparison test; shSbl vs. shAPLP1, p = 0.0385) (Fig. 3D). Therefore, astrocyte APP and APLP2 have little effect on the formation or the stability of excitatory synapses *in vitro* whilst astrocytic APLP1 appears to modulate the extent of postsynapse assembly *in vitro*.

### APLP1 and APLP2 favour astrocyte morphology elaboration

We find that astrocytic APP and APLPs modulate astrocyte morphological elaboration. Because they are cell adhesion molecules capable of homophilic and heterophilic interactions in *trans* (Soba et al., 2005; Schilling et al., 2017), astrocyte morphological complexity could rely on the interaction of astrocytic APP and APLPs with neuronal APP, APLP1 and APLP2. We next sought to test the ability of non-astrocytic APP, APLP1 or APLP2 as a partner in promoting astrocyte process elaboration. To this end we used HEK-astrocyte co-cultures in which astrocytes were grown for 24 h on top of HEK cells overexpressing a candidate adhesion protein partner. Notably, amongst APP isoforms, a previous study reported that APP695 was enriched in neurons whereas APP751 was the prevalent isoform in astrocytes (LeBlanc et al., 1991). Therefore, for APP, we used the 695 isoform for expression in HEK cells. Moreover, neurexin 1β (NRXN1β) was shown to be an effective minimal component in promoting the morphological elaboration of astrocytes, when astrocytes were plated on top of nanofibers coated with NRXN1β (Stogsdill et al., 2017). We thus chose to include NRXN1β as a positive control while we also tested neuroligin 1 (NL1) as a possible non-astrocytic factor in affecting astrocyte morphology. HEK cells expressing mGFP served as a negative control. Astrocytes were labelled with GFAP to reveal the major astrocytic branches, and the complexity of GFAP signals was quantified by a Sholl analysis as described above (Fig. 4A). Astrocytes grown on top of HEK cells did not display the typical fibroblast-like morphology they showed when they were grown alone (Fig. 1B and Fig. 4A). In particular, astrocytes developed more branches when grown on top of HEK cells expressing NRXN1β, APLP1 or APLP2 while neuroligin 1 (NL1) or APP did not seem to affect astrocyte process complexity (N = 7-8 cultures, repeated measures two-way ANOVA with Dunnett’s multiple comparison test at 40 µm distance; mGFP vs. NRXN1β, p = 0.0083; mGFP vs. APLP1, p = 0.0365; mGFP vs. APLP2, p = 0.0215) (Fig. 4B). Thus, APLP1 and APLP2 are potential trans-cellular adhesion partners of non-astrocytic cells such as neurons, which promote process elaboration of astrocytes.

**Figure 4:**
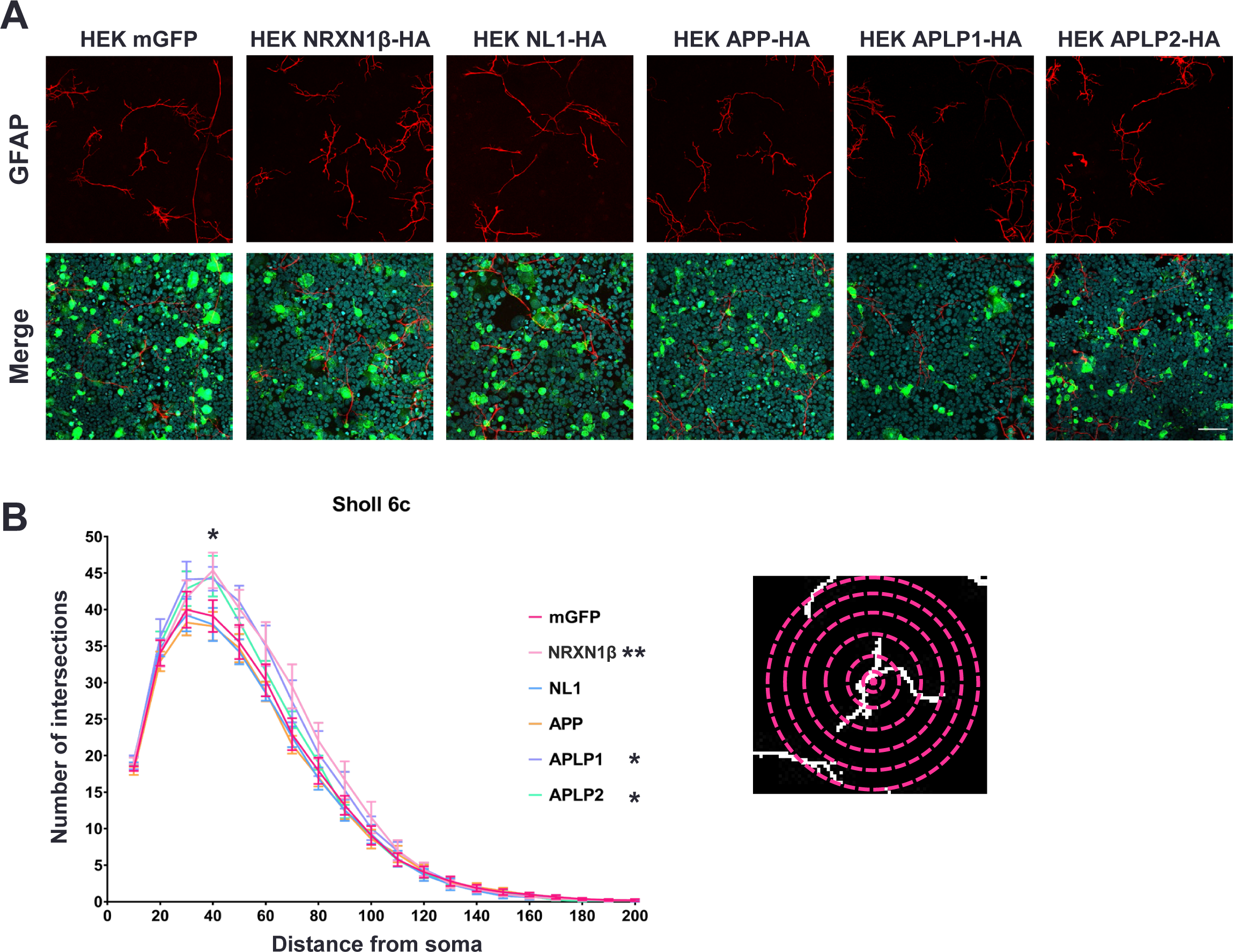
APLP1 and APLP2 promote astrocyte morphology elaboration. **A)** Astrocytes labelled with GFAP (in red) were grown on top of HEK 293 cells overexpressing mGFP, NRXN-1β-HA, NL1-HA, APP-HA, APLP1-HA or APLP2-HA (in green). Images were taken on a confocal microscope with a 20x objective. Scale bar 100µm. **B)** Astrocyte morphology complexity was quantified by Sholl analysis (N = 7-8 cultures, 10 astrocytes per culture, repeated measures two-way ANOVA with Dunnett’s multiple comparison test).

### Astrocytic APP modulates astrocyte morphology *in vivo*

We have observed that astrocytic APP and APLPs can affect astrocyte morphology *in vitro*. However, astrocytes in dissociated cultures do not have the complex and highly ramified morphology found *in vivo*. For this reason, we examined the role of astrocytic APP in regulating astrocyte morphology *in vivo*. We focused on the APP protein because of its well described role in neuronal and synaptic development in general (Dawson et al., 1999; Seabrook et al., 1999; Priller et al., 2006; Bittner et al., 2009; Tyan et al., 2012; Weyer et al., 2014; Martinsson et al., 2019) and in Alzheimer’s disease in the ageing brain, and because we could compellingly knock down its expression in astrocytes *in vivo*. To evaluate the impact of APP KD *in vivo*, AAVs expressing EGFP under the GFAP promoter that also contained the shSbl or shAPP were prepared, in which the expression of shRNAs under a GFAP promoter was enabled by the miR30 sequences flanking the shRNA sequence. AAVs were injected in P0 mice brain using n-SIM method (see Methods). Astrocytic expression of GFP was observed starting as early as P7 and signals remained robust at P14 and P21 (Fig. S5). We adjusted the amount of AAV injected to obtain a sparse astrocyte labelling, and we focused our analyses of the effects of astrocyte APP KD on hippocampus and cortex. In addition, given the role of APP in the developing brain, we confined our analyses to P21 when the bulk of synapse formation had occurred and when astrocytes had been segregated from each other to form their characteristic individual domains and reached a mature complex morphology (Bushong et al., 2004; Saint-Martin and Goda, 2022; Zhong et al., 2023). To confirm astrocytic APP KD *in vivo*, P21 brain sections were processed for immunohistochemistry to label for APP and imaged on a confocal microscope (Fig. S6A). Using the IMARIS surface tool the volume of APP fluorescence signal inside GFP positive astrocytes was evaluated and divided by the total volume of the astrocyte (Fig. S6B). Astrocytic APP expression was significantly decreased in the shAPP condition (N = 6 mice, 1-2 slice per mouse, 5 astrocytes per section; Mann-Whitney test, p = 0.0011) (Fig. S6B). Of note, due to limitations in resolution, we cannot exclude the possibility that part of the selected APP signal might come from neighbouring synapses.

Astrocyte morphology was assessed based on the GFP fluorescence, which allowed the visualization of the entire astrocyte including its small processes (Fig. 5A,C). Neuropil infiltration volume (NIV) is a measure of the volume occupied by astrocyte processes that can vary depending on both astrocyte process number and size, and NIV reflects the complexity of PAPs (Stogsdill et al., 2017). For each astrocyte, three regions away from major branches and cell soma were selected for NIV analysis. We first wondered if astrocytes across hippocampal regions and layers had different NIVs. To investigate this, we used P21 brain sections from mice injected with AAV-pGFAP-EGFP-shEmpty(miR30) allowing astrocytic expression of EGFP without shRNA (Fig. S7A). We found that astrocytes in the *stratum lacunosum moleculare* (SLM) had higher NIVs than in the *stratum radiatum* (SR) or *stratum oriens* (SO) (N = 3 mice, 3 slices/mouse, 12-25 astrocytes/slice, One-way ANOVA followed by Tukey’s multiple comparison test: SR vs. SLM, p = 0.0011; SLM vs. SO, p = 0.0111) (Fig. S5B,C). When comparing astrocyte NIVs between the CA3, CA2 and CA1 regions, no significant differences were found while the trend for a higher NIVs for astrocytes in SLM compared to those in SR and SO persisted in each region (Fig. S7B,C). We further quantified the effects of compromising astrocytic APP expression on NIVs by focusing on CA3 SR. We found that NIVs were decreased in APP KD astrocytes compared to the shSbl control astrocytes in CA3 SR (N = 3 mice, 2 slices/mouse, 5 astrocytes/slice, unpaired t test p = 0.0053) (Fig. 5B). We also examined the impact of APP KD on NIV in astrocytes in the cortex, in the somatosensory area. NIV was significantly decreased in shAPP astrocytes compared to shSbl astrocytes in the layer V/VI where labelled astrocytes were reliably present (N = 5 mice, 1-3 slice/mouse, 5 astrocytes/slice, Mann-Whitney test p = 0.0003) (Fig. 5D)

**Figure 5:**
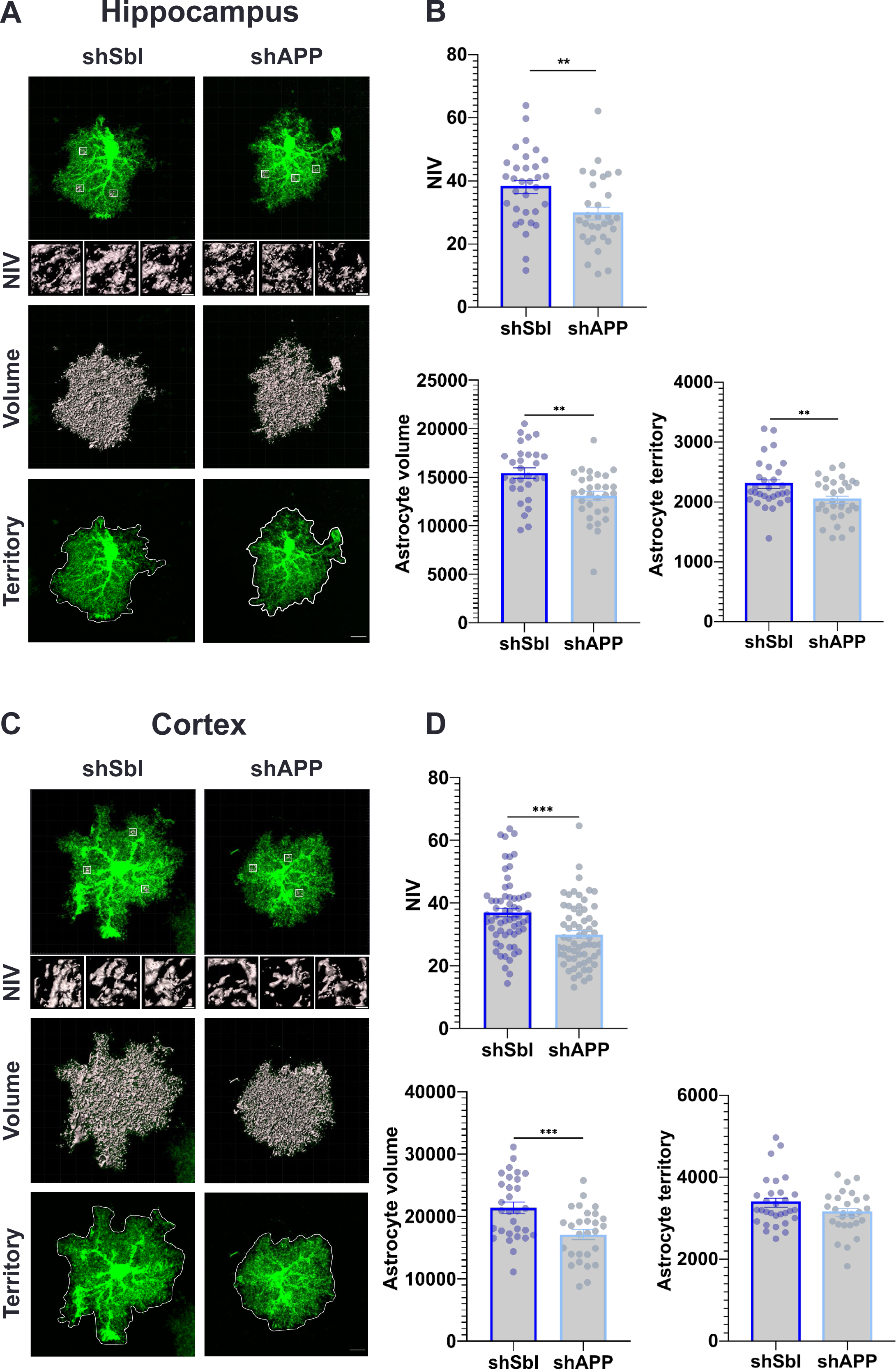
Impact of astrocytic APP on astrocyte morphology elaboration *in vivo*. **A)** Representative images of hippocampus CA3 SR shSbl and shAPP astrocytes, neuropil infiltration volumes (NIVs), total cell volume and territory. Images were taken on a confocal microscope with a 60x silicon immersion objective and a 2x optical zoom. Scale bar 1µm and 10µm. **B)** Quantitative analysis for shSbl and shAPP CA3 SR hippocampal astrocyte NIV (N= 3 mice, 2 slices/mouse, 5 astrocytes/slice, unpaired t test), volume and territory (N= 3 mice, 2 slices/mouse, 5 astrocytes/slice, unpaired t test). **C)** Representative images of somatosensory cortex layer V/VI shSbl and shAPP astrocytes, neuropil infiltration volumes (NIVs), total cell volume and territory. Images were taken on a confocal microscope with a 60x silicon immersion objective and a 2x optical zoom. Scale bar 1µm and 10µm. **D)** Quantitative analysis for shSbl and shAPP somatosensory cortex layer V/VI astrocyte NIV (N= 5 mice, 1-3 slice/mouse, 5 astrocytes/slice, Mann-Whitney test), volume and territory (N= 3 mice, 2 slices/mouse, 5 astrocytes/slice).

We next analysed the volume and territory occupied by astrocytes. To image entire astrocytes, 100 µm thick sections were used. Astrocytes were 3D reconstructed using IMARIS software, and the whole cell volume was measured (Fig. 5A,C). We found that astrocyte volume was decreased in shAPP compared to shSbl astrocytes in the hippocampus and the cortex (N = 3 mice, 2 slices/mouse, 5 astrocytes/slice, unpaired t test: hippocampus, p = 0.0016; cortex, p = 0.0009) (Fig. 5B,D). To measure the 2D territory of astrocytes, we used maximum intensity projections and determined the astrocyte area (Fig. 5A,C). While astrocyte territory was decreased in hippocampal astrocytes, no change was found in the cortex (N = 3 mice, 2 slices/mouse, 5 astrocytes/slice, unpaired t test: hippocampus, p = 0.0077) (Fig. 5B,D).

To confirm the observed effects of APP KD, we sought to rescue the shAPP phenotype by an AAV containing the human APP751 with a C-terminal HA-tag under a GFAP promoter. The APP751 isoform was chosen because of its high expression in astrocytes (LeBlanc et al., 1991), and moreover, the shAPP sequence was not designed to knockdown the human APP (Young-Pearse et al., 2007). For this experiment, shSbl or shAPP AAVs were injected either alone or co-injected with APP751-Ct-HA AAV (Fig. 6A, Fig. S6A). APP expression level in astrocytes was observed by immunofluorescence labelling using an antibody against APP on brain sections (Fig. S8A). Quantification of the APP signal inside transduced astrocytes expressing GFP indicated a 3-4 fold increase in APP expression by the rescue AAV (Fig. S8B). To expedite the identification of APP751-Ct-HA expressing astrocytes, an antibody against the HA-tag was used (Fig. 6A). Measurements of NIVs in each condition in cortical astrocytes (somatosensory cortex layer V/VI) showed that the decrease in NIV found in shAPP astrocytes was rescued by the expression of APP751 (N = 2 mice, 3 slices/mouse, 5 astrocytes/slice, One-way ANOVA followed by Tukey’s multiple comparison test: shAPP vs. shSbl, p = 0.0051; shAPP vs. shAPP+APP, p = 0.0364) (Fig. 6B). Interestingly, APP751 overexpression in shSbl astrocytes did not further increase NIV compared to NIV in control shSbl alone astrocytes (Fig. 6B). These results indicate that astrocytic APP, when knocked down, can specifically reduce astrocyte process elaboration. In the cortex, APP KD might lead to a decrease in process density without impacting their length, such that defining of the astrocyte territory borders may involve a process that is less dependent on the levels of APP. In addition, the finding that overexpression of APP751 in control astrocytes does not increase NIV suggests that astrocytic APP may not be sufficient on its own to promote astrocyte morphological complexity and that there may be a ceiling for astrocytic APP level for its function in facilitating astrocyte process elaboration, for example, by limited availability of partner molecules.

**Figure 6:**
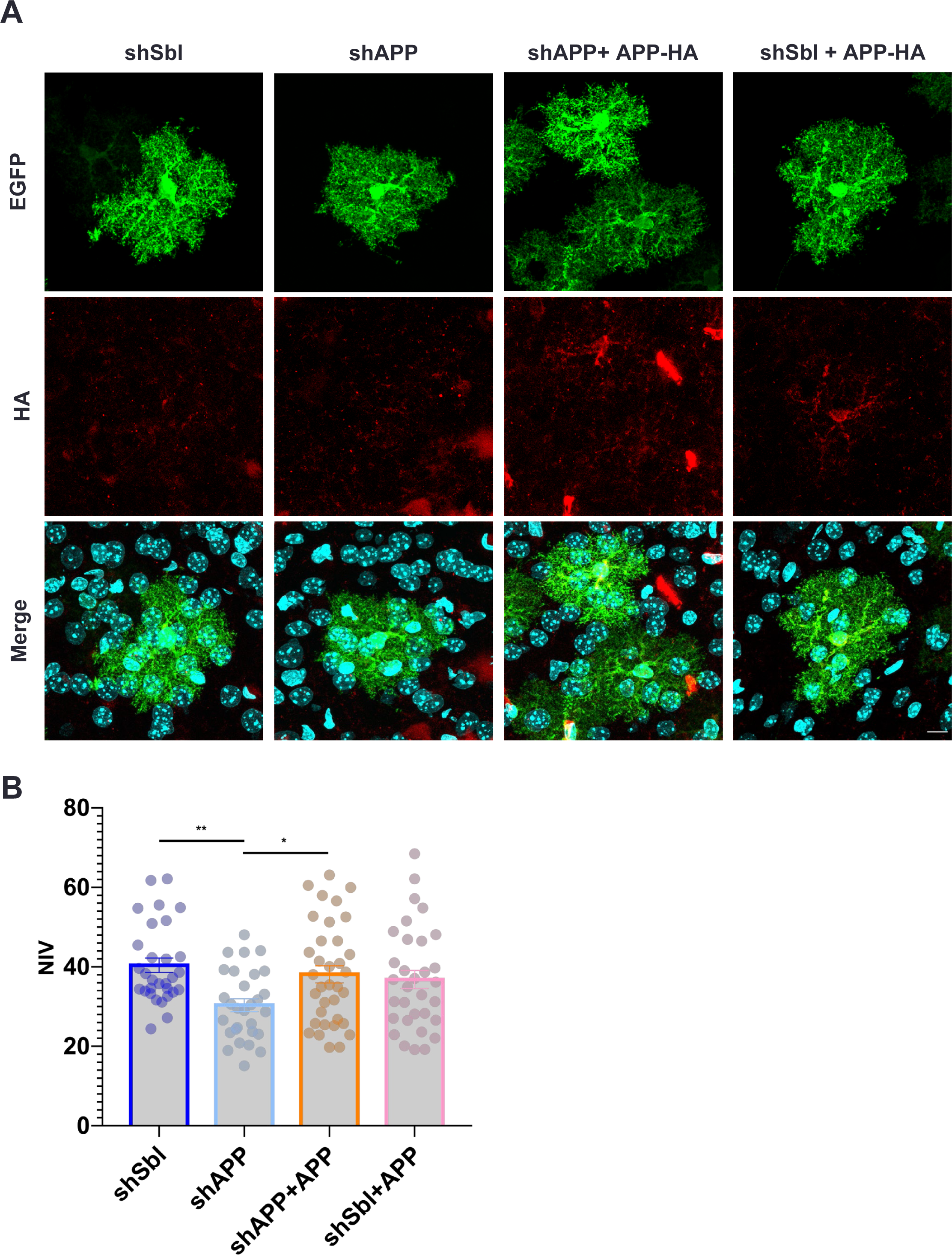
Impact of APP overexpression or APP rescue on NIV *in vivo*. **A)** Somatosensory cortex layer V/VI astrocytes expressing EGFP (green) with shSbl or shAPP alone or with APP751-Ct-HA labelled with anti-HA antibodies (red). Scale bar 10µm. **B)** Quantitative analysis for shSbl, shAPP, shAPP+APP751 (rescue) and shSbl+APP751 cortical astrocyte NIV (N= 2 mice, 3 slices/mouse, 5 astrocytes/slice, One-way ANOVA followed by Turkey’s multiple comparison test).

### Astrocytic APP has a limited impact on synapses *in vivo*

*In vivo* studies have reported synaptic alterations in APP KO mice (Seabrook et al., 1999; Priller et al., 2006; Bittner et al., 2009; Tyan et al., 2012; Martinsson et al., 2019). We therefore asked whether astrocytic KD of APP could also impact synapses *in vivo*. To this end, 20 µm thick brain sections from mice injected with AAVs expressing EGFP and shSbl or shAPP under the GFAP promoter were examined by immunohistochemistry. Excitatory synapses were labelled with antibodies against the presynaptic VGLUT1 and postsynaptic Homer1 (Fig. 7A), and inhibitory synapses were assessed using antibodies against presynaptic VGAT and postsynaptic Gephyrin (Fig. 7C). Two optical planes containing an astrocyte from the cortex (somatosensory layer V/VI) were imaged with a confocal microscope and maximum intensity projections were obtained. The sparse astrocyte transduction allowed us to directly compare synapse densities between the area occupied by the transduced astrocyte and the area represented by the non-transduced wild type astrocytes within each sample, to limit technical variability between slices. The number of presynaptic (VGLUT1 or VGAT), postsynaptic (Homer1 or Gephyrin) and co-localized (VGLUT1/Homer1 or VGAT/Gephyrin) puncta inside and outside transduced astrocyte was determined, and for each image, the ratio of number of puncta inside/outside the target transduced astrocyte (in/out) was calculated. For the excitatory synapses we found that the in/out ratio of Homer1 puncta was increased in shAPP astrocytes compared to shSbl controls (N = 3 mice, 2 slice/mouse, 5 astrocyte/slice, unpaired t-test: p = 0.0181) (Fig. 7B). The number of VGLUT1 puncta or co-localized puncta remained unchanged (Fig. 7B). For inhibitory synapses, no differences in gephyrin, VGAT or co-localised puncta were observed between shSbl and shAPP astrocytes (Fig. 7D). Overall, loss of astrocytic APP resulted in an apparent increase in the density of postsynaptic scaffold, Homer1, but did not affect the density of other synaptic markers. Therefore, astrocytic APP might in general have a limited impact on synapses during development while serving to match postsynaptic scaffold assembly to presynaptic compartments at excitatory synapses.

**Figure 7:**
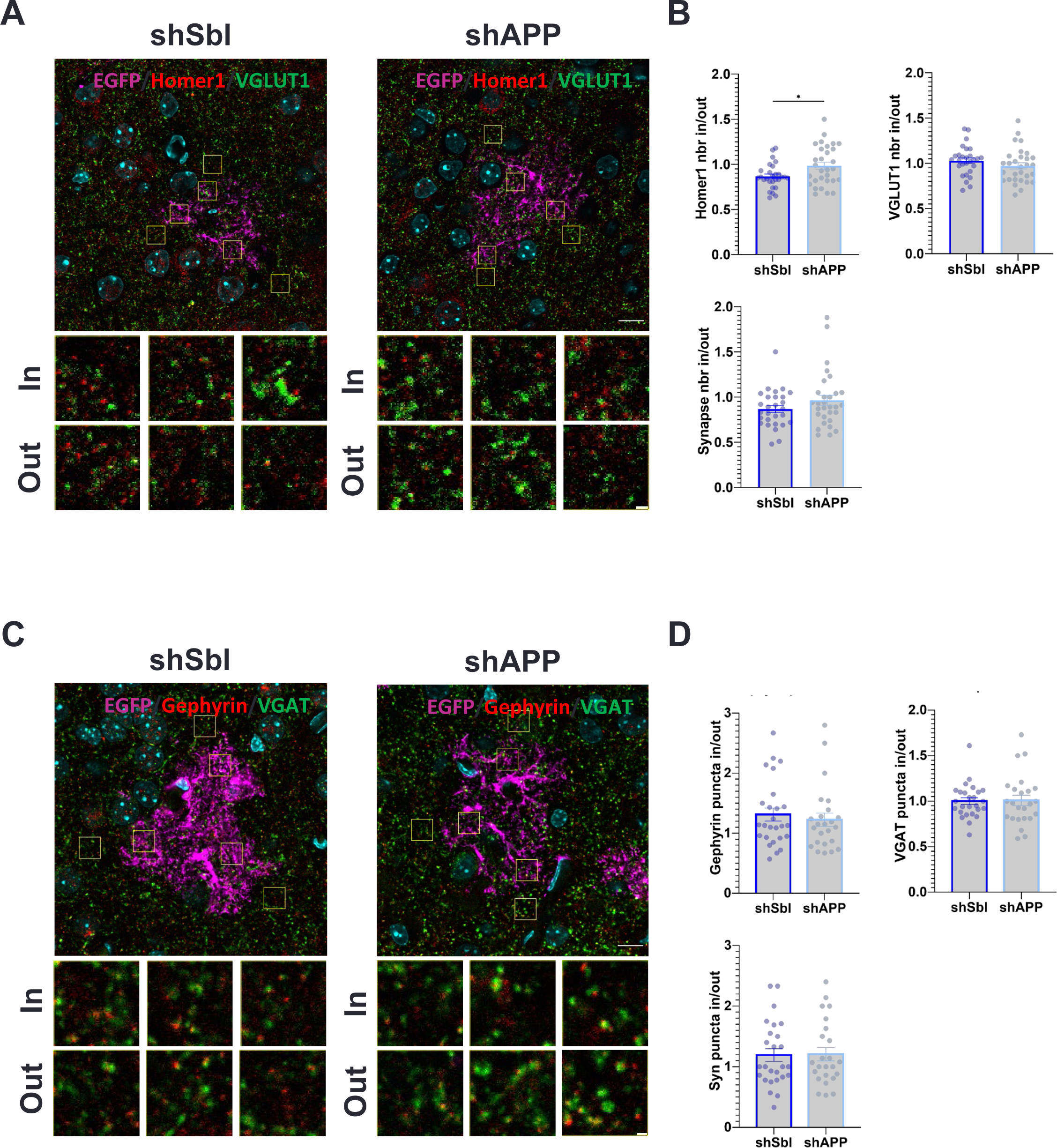
APP KD has a limited impact on synapse formation *in vivo*. **A)** Representative images of somatosensory cortex layer V/VI shSbl and shAPP astrocytes (purple) with excitatory post-synaptic Homer1 (red) and pre-synaptic VGLUT1 (green) labelling. Images were taken on a confocal microscope with a 60x silicon immersion objective and a 2x optical zoom. Three regions inside and outside the EGFP positive astrocyte were selected for analysis with the puncta analyser plugin. Scale bar 1µm and 10µm. **B)** Quantitative analysis of Homer1, VGLUT1 and co-localized (synapses) puncta inside and outside (in/out ratio) astrocyte area (N= 3 mice, 2 slice/mouse, 5 astrocyte/slice, unpaired t-test). **C)** Representative images of somatosensory cortex layer V/VI shSbl and shAPP astrocytes (purple) with inhibitory post-synaptic Gephyrin (red) and pre-synaptic VGAT (green) labelling. Images were taken on a confocal microscope with a 60x silicon immersion objective and a 2x optical zoom. Three regions inside and outside the EGFP positive astrocyte were selected for analysis with the puncta analyser plugin. Scale bar 1µm and 10µm. **D)** Quantitative analysis of Gephyrin, VGAT and co-localized (synapses) puncta inside and outside (in/out ratio) astrocyte area (N= 3 mice, 1-2 slice/mouse, 5 astrocyte/slice, unpaired t test)

## Discussion

### APP and APLPs are expressed in astrocytes

Recent studies have highlighted the expression of synaptic cell adhesion molecules in astrocytes such as neuroligins or ephrins that have traditionally been studied for their roles in bridging presynaptic and postsynaptic neurons (Tan and Eroglu, 2021; Saint-Martin and Goda, 2022). APP and its family members, APLP1 and APLP2 are expressed in the presynaptic and postsynaptic compartments of neurons, where they can function as synapse adhesion molecules (Schilling et al., 2017). However, their expression in astrocytes have been controversial, and if present, the precise roles of astrocytic APP, APLP1 and APLP2 have not been clear. Given the close proximity between synapses and astrocyte processes that are of very small size, the presence of synaptic molecules in astrocytes can be challenging to assess. Supported by the publicly available transcriptomic and translatomic datasets, here we have demonstrated the astrocytic protein expression of APP, APLP1 and APLP2 by combining western blot and immunofluorescence labelling analyses. The bioinformatic datasets represented astrocyte gene expression obtained by cell purification/isolation or by driving astrocyte-specific expression to tag ribosomal proteins and their bound mRNA (TRAP). The presence of App, Aplp1 and Aplp2 mRNA in astrocytes was confirmed in all the datasets examined. Importantly, App, Aplp1 and Aplp2 mRNA were found in astrocyte PAPs (Sakers et al., 2017; Mazaré et al., 2020), which ideally positioned them to be involved in astrocyte-synapse interactions. Of note, actively translated App mRNA seemed particularly enriched in PAPs compared to the levels found in the whole astrocyte (Sakers et al., 2017). This result could explain the difficulties in finding APP signal in astrocytes in immunolabelling experiments based on signal co-localisation with GFAP to identify astrocytes, as GFAP does not reveal small astrocyte processes. In addition, our western blot and immunolabelling analysis indicated that APP, APLP1 and APLP2 proteins were expressed in astrocytes at lower levels than in neurons. This was also the case for transcript level as App mRNA was consistently detected at a lower level in astrocytes compared to neurons or the whole cortical lysates in the published datasets. Interestingly, App, Aplp1 or Aplp2 mRNA were present at a similar or higher level in astrocytes than Nlgn2, a synaptic cell adhesion protein involved in astrocyte-synapse interactions (Stogsdill et al., 2017). Therefore, even a low level of expression of APP and APLPs might suffice to mediate astrocytic influence on synapses.

### APP and APLPs impact astrocyte process elaboration *in vitro*

We have shown that KD of astrocytic APP or APLPs impacts astrocyte process elaboration in culture. Interestingly, while APP and APLP1 KD both decreased the number of astrocytic processes relative to the control shSbl condition, APLP2 KD had the opposite effect in promoting astrocyte process complexity. In addition, decreases in the maximum process length and the total area of astrocytes were observed only for APLP1 KD astrocytes and not for APP KD astrocytes. Collectively, these differences in the effects of compromising astrocytic expression of APP, APLP1 and APLP2 on astrocyte process elaboration are reminiscent of the differential phenotypes observed in APP and APLP1 KO mice models (Tyan et al., 2012; Schilling et al., 2017).

Astrocytic APP, APLP1 and APLP2 could mediate astrocyte process elaboration by interacting with distinct set of molecules that are expressed in neurons. Notably, when exogenously overexpressed in HEK cells as non-astrocytic interacting substrates, we find that similarly to NRXN-1β, APLP1 and APLP2 are capable of promoting astrocyte process elaboration. Therefore, it is possible that astrocytic APP or APLP1 could interact with neuronal APLP1 or APLP2 to promote astrocyte process elaboration. Interestingly, NRXN-1β has been recently shown to interact with APP in *trans* (Cvetkovska et al., 2022). NRXN-1β that is expressed in the presynaptic compartment could thus act as a neuronal partner of astrocytic APP to facilitate the formation and/or maintenance of complex astrocyte processes, although the mechanistic basis by which such adhesive interactions promotes process branching warrant further studies.

### APP and APLP1 KD in astrocytes have a limited impact on synapse density

To evaluate the effect of knocking down astrocytic APP or APLPs on synapse formation and/or stability we have labelled excitatory or inhibitory synapses in our *in vitro* and *in vivo* models. In rat hippocampal neuron-astrocyte co-cultures, compromising astrocytic expression of APLP1 increases Homer1 puncta density without affecting VGLUT1 puncta or synaptic density (i.e. density of co-apposed VGLUT1/Homer puncta), which suggests that the loss of APLP1 facilitates the clustering of Homer1 scaffold. In contrast, we find no significant effect of astrocytic APP or APLP2 KD on synapse formation in cultures. However, *in vitro* models have limitations in that cultured astrocytes do not acquire the mature complex morphology they develop *in vivo*. Moreover, astrocytes in our cultures show highly overlapping processes and do not become segregated from each other as they do *in vivo*. When APP is sparsely knocked down in astrocytes *in vivo*, we could observe a slight increase in Homer1 puncta density. Therefore, astrocytic APP *in vivo* might help limit synaptic Homer1 scaffolds, akin to the finding for APLP1 *in vitro*. No alteration is observed in inhibitory synapse density, and thus a decrease in APP expression in astrocyte has overall a limited impact on synapse formation and/or stability. It is however possible that stronger effects on synapses and astrocyte processes might be observed with a complete knockout of astrocytic APP or upon knocking out both astrocytic APP and APLP2 as APLP2 has been suggested to have redundant functions with APP (Heber et al., 2000). In addition, to facilitate identification of NIVs in target astrocytes, astrocytes in our models were sparsely knocked down for APP; however, a loss of APP uniformly in all astrocytes might lead to a different phenotype. Interestingly, in previous reports, despite the conflicting evidence concerning the effects of APP KO on synapses, the synaptic effects *in vivo* when observed, seemed to be age dependent. For example, presynapse or postsynapse density was decreased in aged APP KO mice (Seabrook et al., 1999; Lee et al., 2010; Tyan et al., 2012) but conversely increased in young or young-adult APP KO mice (Priller et al., 2006; Bittner et al., 2009; Martinsson et al., 2019). Therefore, these findings suggest that APP’s functions and/or its mode of action might differ with the age of animal. Of note, the increased excitatory postsynaptic marker density within APP KD astrocyte territory in our model is in line with these observations. Further mechanistic insights into how astrocytic APP regulates synapse formation and/or stability, specifically, the apposition of presynaptic and postsynaptic compartments, would be valuable in understanding the age-dependent effects of global (i.e. both neuronal and astrocytic) APP loss on presynaptic and postsynaptic components.

### APP KD *in vivo* decreases astrocyte volume and neuropil infiltration

In addition to a decrease in astrocyte process elaboration in APP KD astrocytes *in vitro*, we showed that APP KD *in vivo* decreased astrocyte neuropil infiltration and total volume in both the cortex and the hippocampus. Importantly, APP751 overexpression could rescue the decrease in NIV observed in APP KD cortical astrocytes. Of note, APP751 overexpression *per se* does not appear to increase astrocyte NIV. In one possible explanation, the space in neuropil could be already at capacity to accommodate astrocyte processes such that further growth is not feasible. In another possibility, a lack of effect of overexpression of APP751 could indicate additional mechanisms at play for APP-dependent increase in astrocyte NIVs that involves, for instance, other interacting APP partners. Overall, our results point towards a role of APP in development of astrocyte morphological complexity, especially the formation of new processes. Whether this function relies solely on full length APP or involves its various cleavage products remains to be investigated. APP-derived fragments have been shown to impact neuronal functions (Nhan et al., 2015). For example, in neurite outgrowth assays, secreted, soluble form of APP, APPsα, is thought to impair neurite lengthening by hindering the interaction between full length APP and integrin-β1 (Young-Pearse et al., 2008). Furthermore, APPsα has been reported also to exert neuroprotective and synaptogenic activities (Nhan et al., 2015); in APP KO and APP/APLP2 double KO mice, APPsα rescues the deficits in LTP and dendritic spine density (Ring et al., 2007; Weyer et al., 2014; Richter et al., 2018). Interestingly, a recent study has shown that APP695sα modulates the proteome and secretome of mouse primary astrocytes with changes observed in proteins involved in cell morphology and vesicle dynamics (Peppercorn et al., 2023). Additionally, APP intracellular domain (AICD) can modulate transcriptional activity and induce the expression of genes involved in the organisation and dynamics of actin in human neural cell cultures (Müller et al., 2007). Altogether, the contribution of astrocytically derived fragments of APP in regulating astrocyte morphology and astrocyte-synaptic interactions warrants further investigation.

In considering the literature for APP, most prior studies have addressed neuronal APP and its effects on neurons, which mainly express the APP695 isoform. Unlike neurons, astrocytes predominantly express APP751 and 770 isoforms that contain a Kunitz protease inhibitor (KPI) domain, which is absent in APP695. Analysis of the levels of each App mRNA isoform in rat cortical astrocyte cultures reveals distinct temporal expression patterns (LeBlanc et al., 1991). APP751 mRNA levels remain stable between DIV0 and DIV21 whilst APP695 strongly decreases and APP770 increases (LeBlanc et al., 1991). With such differences in timing and the prevalent cell type of expression, it is possible that APP isoforms serve distinct functions. In favour of this hypothesis, APP695 and APP751 show differences in their interactomes in SH-SY5Y neuronal cells (Andrew et al., 2019). Consistently with such differences, APP751 and APP770 isoforms are more potent in promoting neurite outgrowth than APP695 (Qiu et al., 1995), yet they show similar capacities in mediating cell aggregation via *trans* interactions (Isbert et al., 2012). As for their processing, APP695 appears to produce more Aβ42 and nuclear AICD than APP751 and APP770 in SH-SY5Y cells (Belyaev et al., 2010). Other studies suggest an impact of the KPI domain on APP770 and APP751 dimerization: KPI domain of APP770 might impede its *cis* dimerization in the endoplasmic reticulum (Isbert et al., 2012) while another study found an increased dimerization of APP751 compared to APP695 (Khalifa et al., 2012). In summary, how APP isoforms differ in their function remains largely unknown and warrants further investigation, particularly in understanding the specific roles for APP in neurons and glial cells.

## Acknowledgements

This study was carried out while authors were affiliated at RIKEN Center for Brain Science. MSM was a recipient of Overseas Research Fellowship of the Japan Society of the Promotion of Science (P19728). The work was supported by RIKEN Center for Brain Science, Okinawa Institute of Science and Technology Graduate University, and the Japan Society for the Promotion of Science Core-to-Core Program (JPJSCCA20170008).

## Supplementary Materials

### Supplemental Figure Legends

**Supplemental Figure 1:**
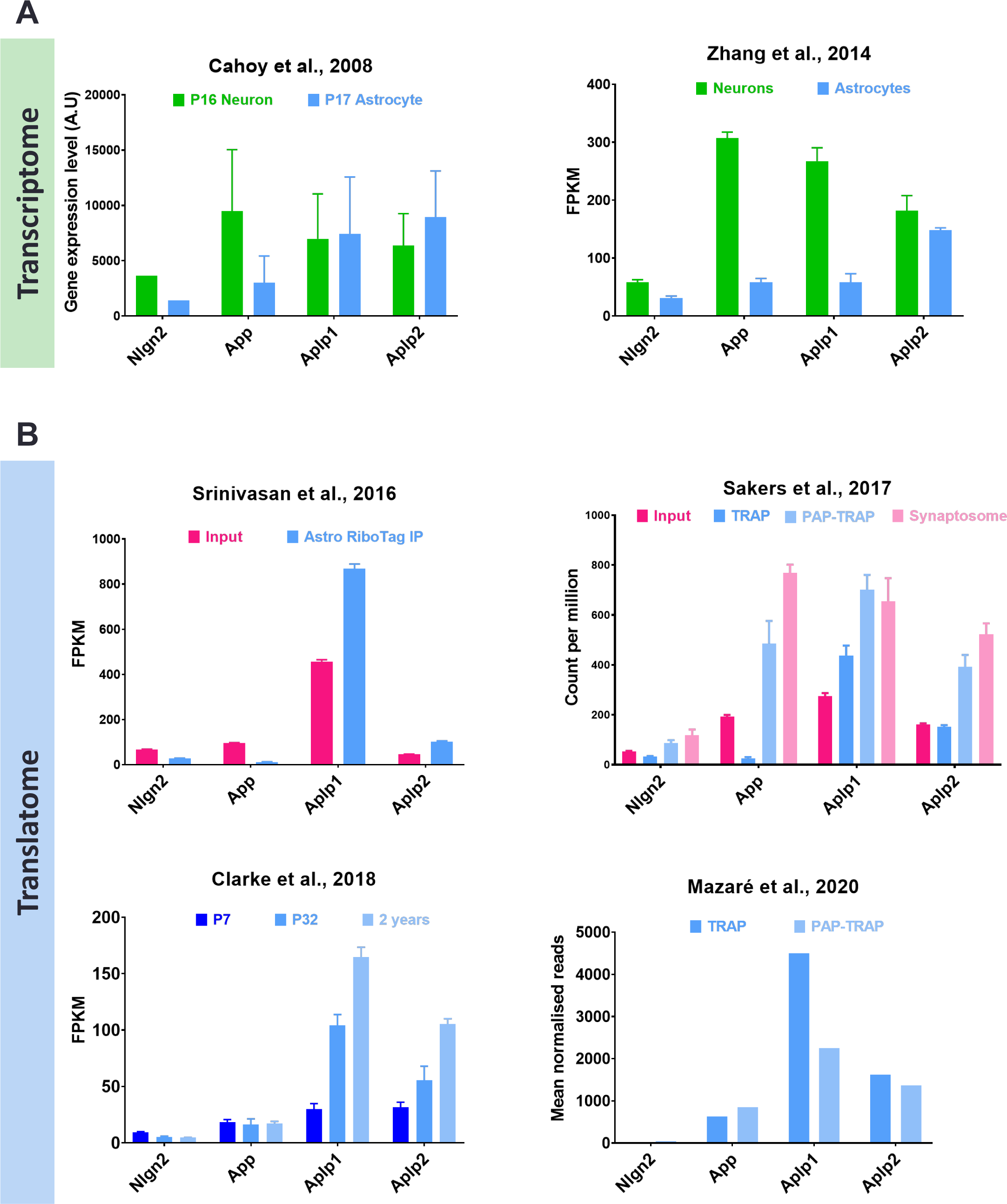
APP and APLPs mRNA are expressed in neurons and astrocytes. **A)** Transcriptome datasets for Nlgn2, App, Aplp1 and Aplp2 mRNA expression in purified neurons (green) and astrocytes (blue) obtained by GeneChip array (Cahoy et al., 2008) and RNA sequencing (Zhang et al., 2014). **B)** Translatome datasets for Nlgn2, App, Aplp1 and Aplp2 mRNA in astrocytes (blue) from the cortex (Srinivasan et al., 2016; Sakers et al., 2017) and the hippocampus (Clarke et al., 2018; Mazaré et al., 2020). Actively translated mRNA from astrocytes was isolated by translating ribosome affinity purification (TRAP) using mice with astrocyte specific expression of tagged ribosomal proteins. Actively translated mRNA from perisynaptic astrocyte processes (PAPs) was obtained by performing a TRAP experiment on purified synaptosomes (PAP-TRAP) (Saker et al., 2017; Mazaré et al., 2020). Actively translated mRNA from hippocampal astrocytes at different time points (P7, P32 and 2years) (Clarke et al., 2018). FPKM = fragments per kilobase of transcript per million reads of mRNA.

**Supplemental Figure 2:**
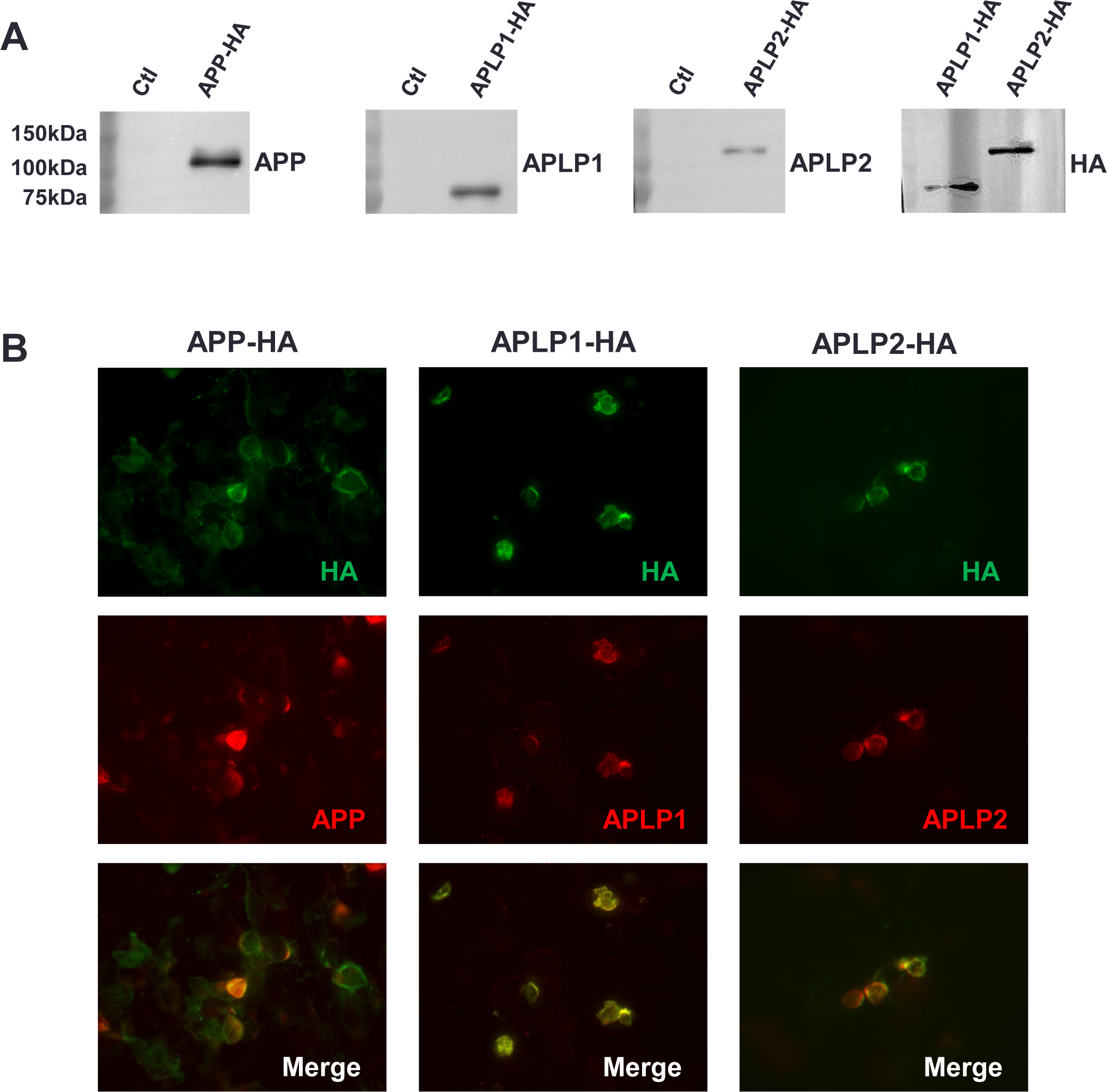
Validation of anti-APP, anti-APLP1 and anti-APLP2 antibodies. **A)** HEK 293 cells transfected to express APP-HA, APLP1-HA or APLP2-HA or non-transfected HEK 293 cells (Ctl) were lysed and analysed by western blot with antibodies against APP, APLP1, APLP2 or with antibodies against the HA-tag. APP, APLP1 and APLP2 bands were found around 110kDa, 80kDa and 120kDa respectively. **B)** HEK 293 cells transfected to express APP-HA, APLP1-HA or APLP2-HA were labelled with anti-APP, APLP1 or APLP2 antibodies respectively and anti-HA-tag antibodies.

**Supplemental Figure 3:**
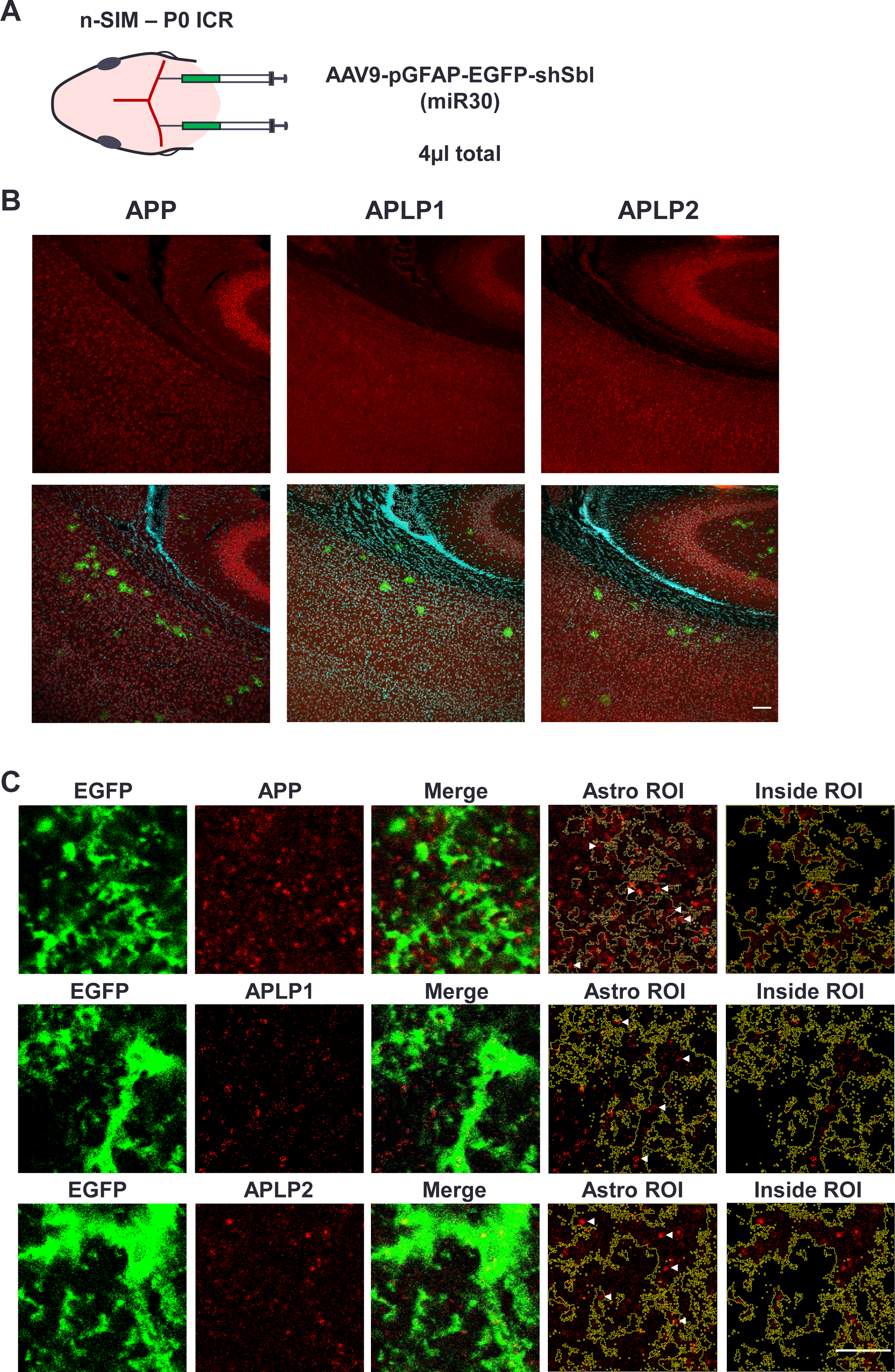
APP, APLP1 and APLP2 expression *in vivo*. **A)** Neonatal sinus injection method. AAV with EGFP and the scramble shRNA under the GFAP promoter for astrocyte specific expression was injected in newborn ICR mice at P0. **B)** Cortex from a P21 mouse injected with AAV9-pGFAP-EGFP-shSbl(miR30) and labelled with antibodies against APP, APLP1 or APLP2. Images were taken on a confocal microscope with a 10x objective. Sparse astrocytic EGFP expression can be observed. Scale bar 100µm. **C)** Localisation of APP, APLP1 and APLP2 in astrocyte processes. Cortical astrocytes were imaged on a confocal microscope with a 60x silicon immersion objective and a 2x optical zoom, only one optical plane is presented. The EGFP expressed in astrocytes allows the visualization of small astrocyte processes. APP, APLP1 and APLP2 puncta can be observed inside (arrowheads) and outside the astrocyte ROI. For visualisation APP signal outside the astrocyte ROI was deleted (Inside ROI image). Scale bar 5µm.

**Supplemental Figure 4:**
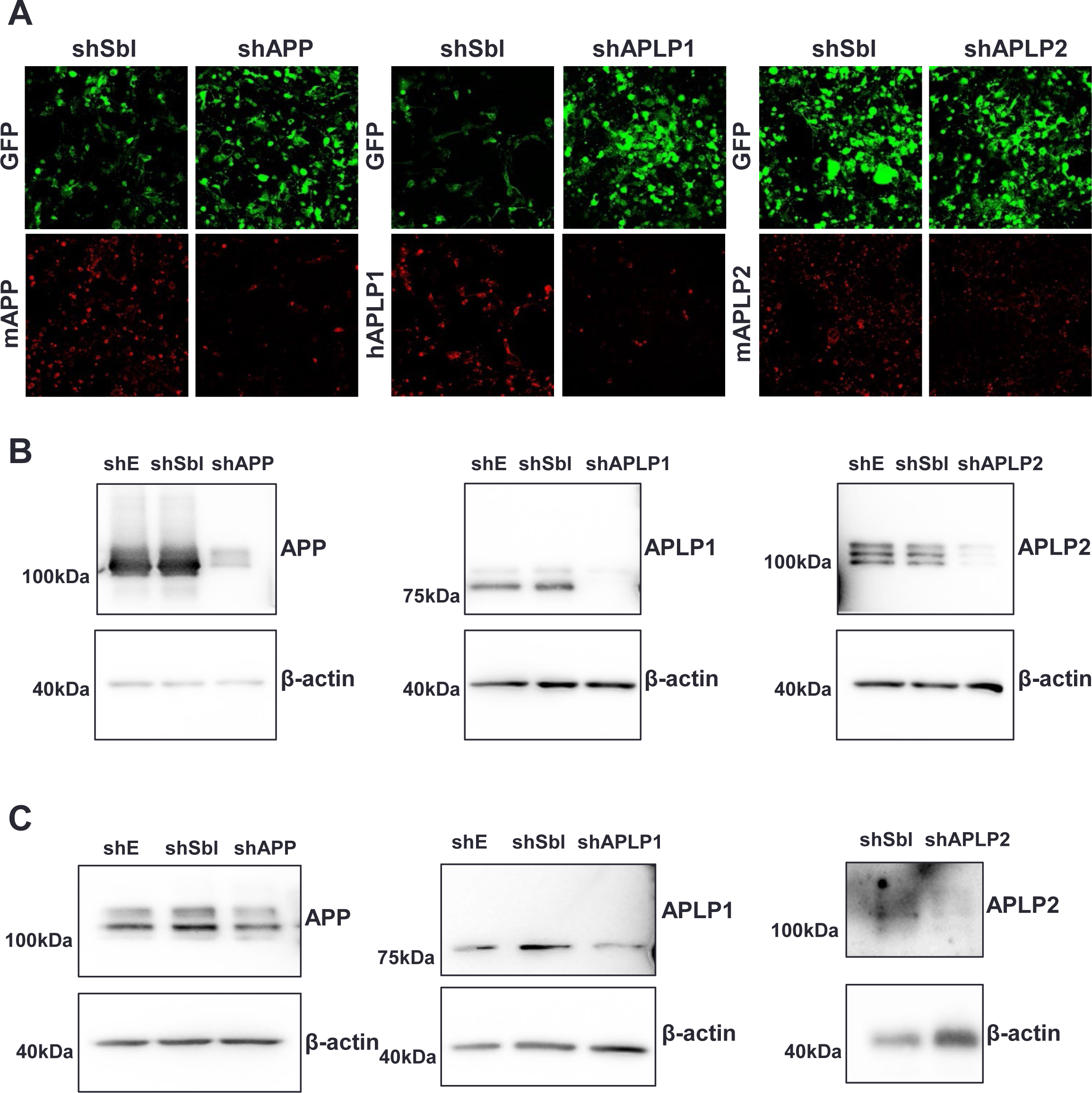
Validation of APP, APLP1 and APLP2 shRNA in HEK cells and rat astrocytes. **A)** HEK cells were transfected with GFP and APP-HA, APLP1-HA or APLP2-HA and the control scramble shRNA (shSbl) or their respective shRNA, shAPP, shAPLP1 or shAPLP2. Cells were then labelled with anti-APP, anti-APLP1 or anti-APLP2 antibodies to visualize protein expression. **B)** HEK cells were transfected with GFP and APP-HA, APLP1-HA or APLP2-HA and the control empty shRNA (shE), scramble shRNA (shSbl) or their respective shRNA, shAPP, shAPLP1 or shAPLP2. Cells were then lysed and APP, APLP1 or APLP2 expression was analysed by western blot with β-actin as a loading control. **C)** Astrocytes were transfected with control empty shRNA (shE), scramble shRNA (shSbl) or with shAPP, shAPLP1 or shAPLP2. Cells were then lysed and APP, APLP1 or APLP2 expression was analysed by western blot with β-actin as a loading control.

**Supplemental Figure 5:**
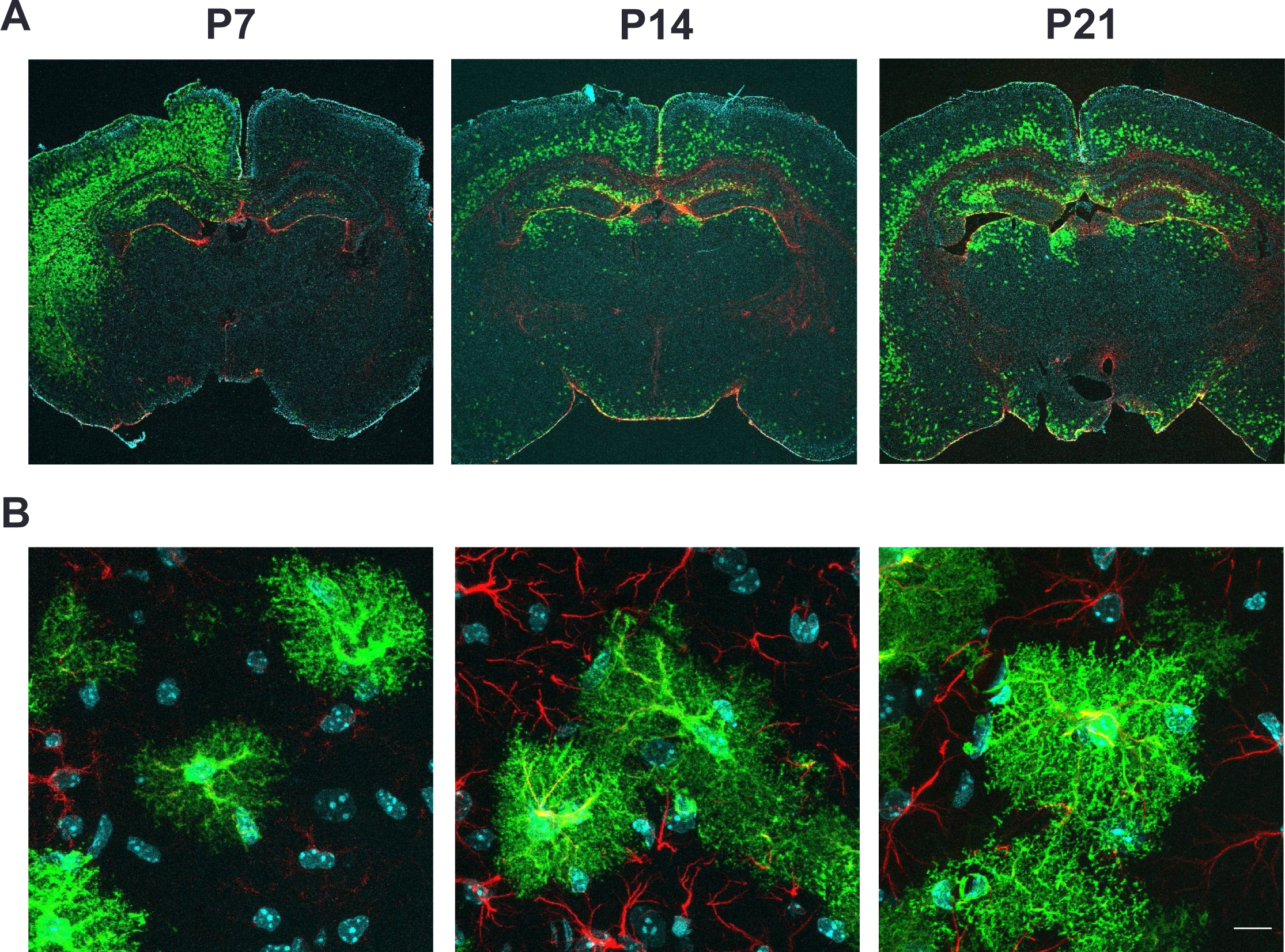
Expression of AAVs in mice at P7, P14 and P21. P0 mice were injected with AAV-pGFAP-EGFP-shE(miR30) and brains were fixed at different ages after injection. **A)** Expression of EGFP is observed in mice cortex and hippocampus at P7, P14 and P21. Images were taken on a confocal microscope with a x2 objective **B)** EGFP expression in astrocytes allows the visualisation of small astrocyte processes. Images were taken on a confocal microscope with a 60x silicon immersion objective and a 2x optical zoom. Scale bar 10µm.

**Supplemental Figure 6:**
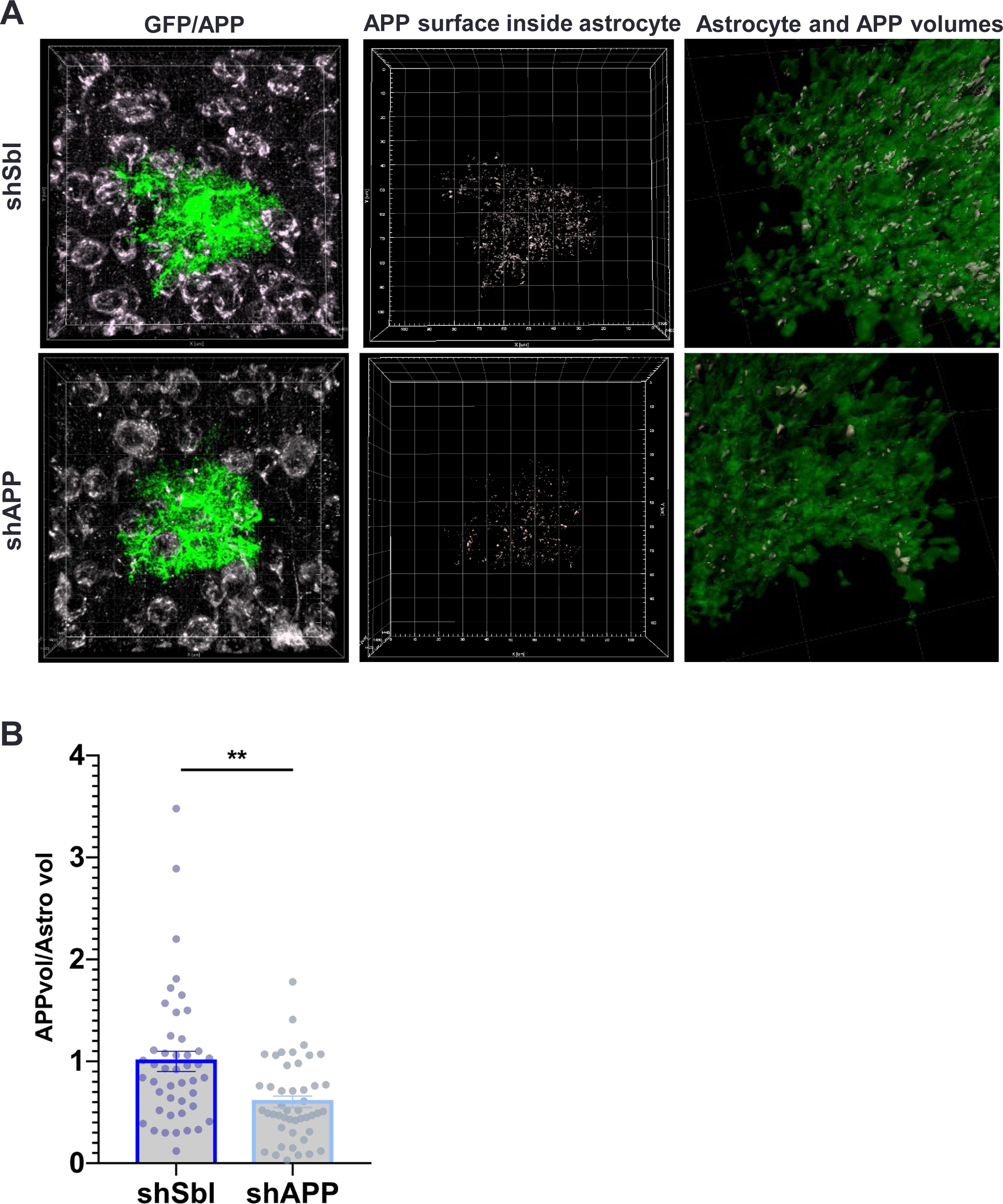
Validation of APP KD *in vivo*. **A)** Brain slices from mice injected with AAV-pGFAP-EGFP-shSbl(miR30) or AAV-pGFAP-EGFP-shAPP(miR30) were labelled with anti-APP antibodies. The volume of APP signal inside EGFP positive astrocytes was reconstructed and measured on IMARIS in shSbl and shAPP astrocytes. **B)** Quantification of the volume of APP (APP vol) per astrocyte volume (Astro vol) in shSbl and shAPP astrocytes normalised to the shSbl condition for each experiment (N=6 mice, 1-2 slice per mouse, 5 astrocyte per slice, Mann-Whitney test).

**Supplemental Figure 7:**
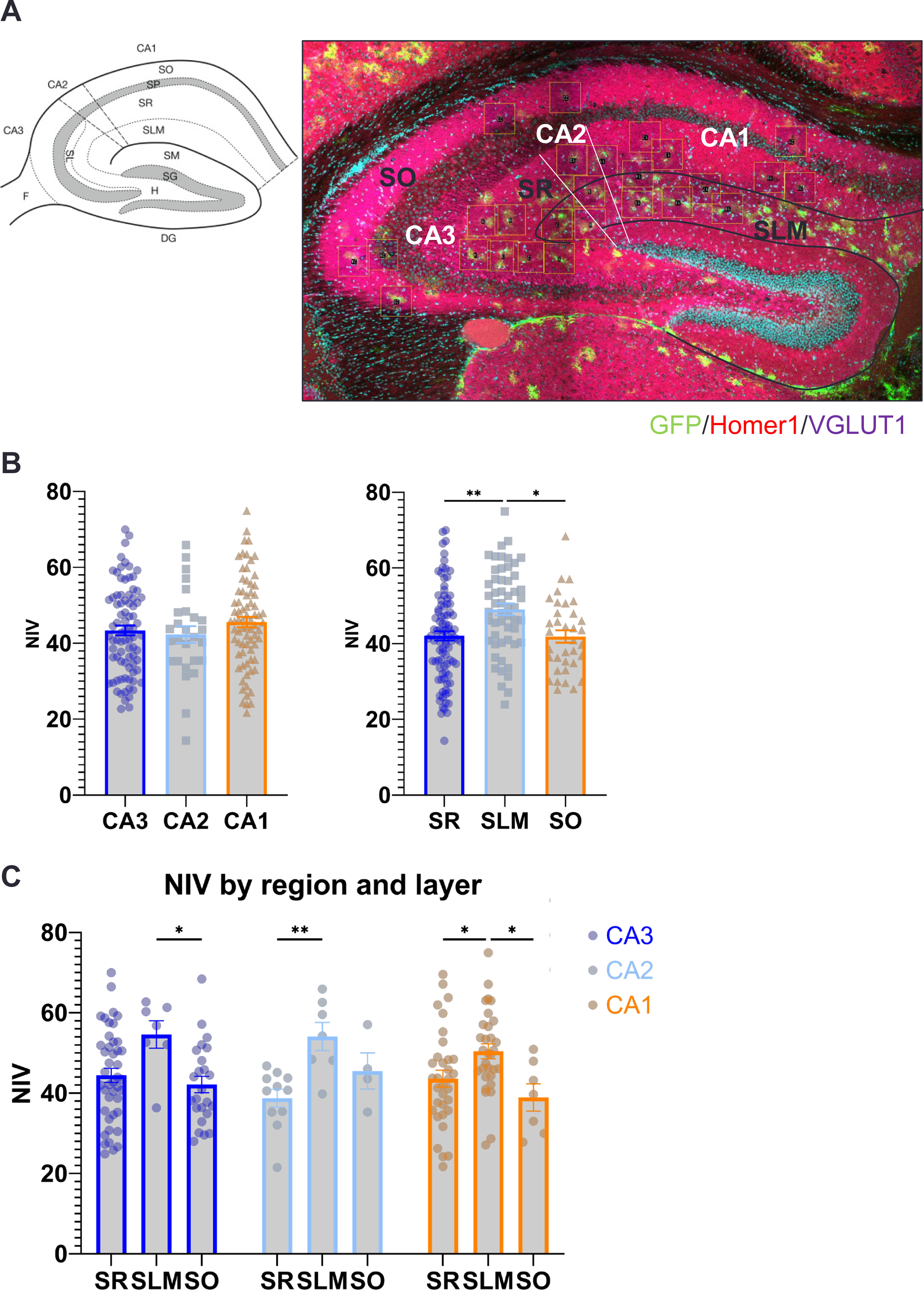
Hippocampus astrocyte NIV in different layers and regions. **A)** Representation of an hippocampus with the different layers and regions and confocal image from a P21 mice hippocampus injected with AAV-EGFP-shE(miR30) and stained for Homer1 and VGLUT1. The imaged astrocytes are localized by squares. **B)** Quantitative analysis of NIVs by regions or layers of the hippocampus (3 slices/mouse, 12-25 astrocytes/slice, one-way ANOVA followed by Turkey’s multiple comparison test). **C)** Detail of astrocyte NIVs in different hippocampal regions depending on the layer. Data were obtained from 3 mice, 3 slices/mouse, 12-25 astrocytes/slice, One-way ANOVA followed by Turkey’s multiple comparison test.

**Supplemental Figure 8:**
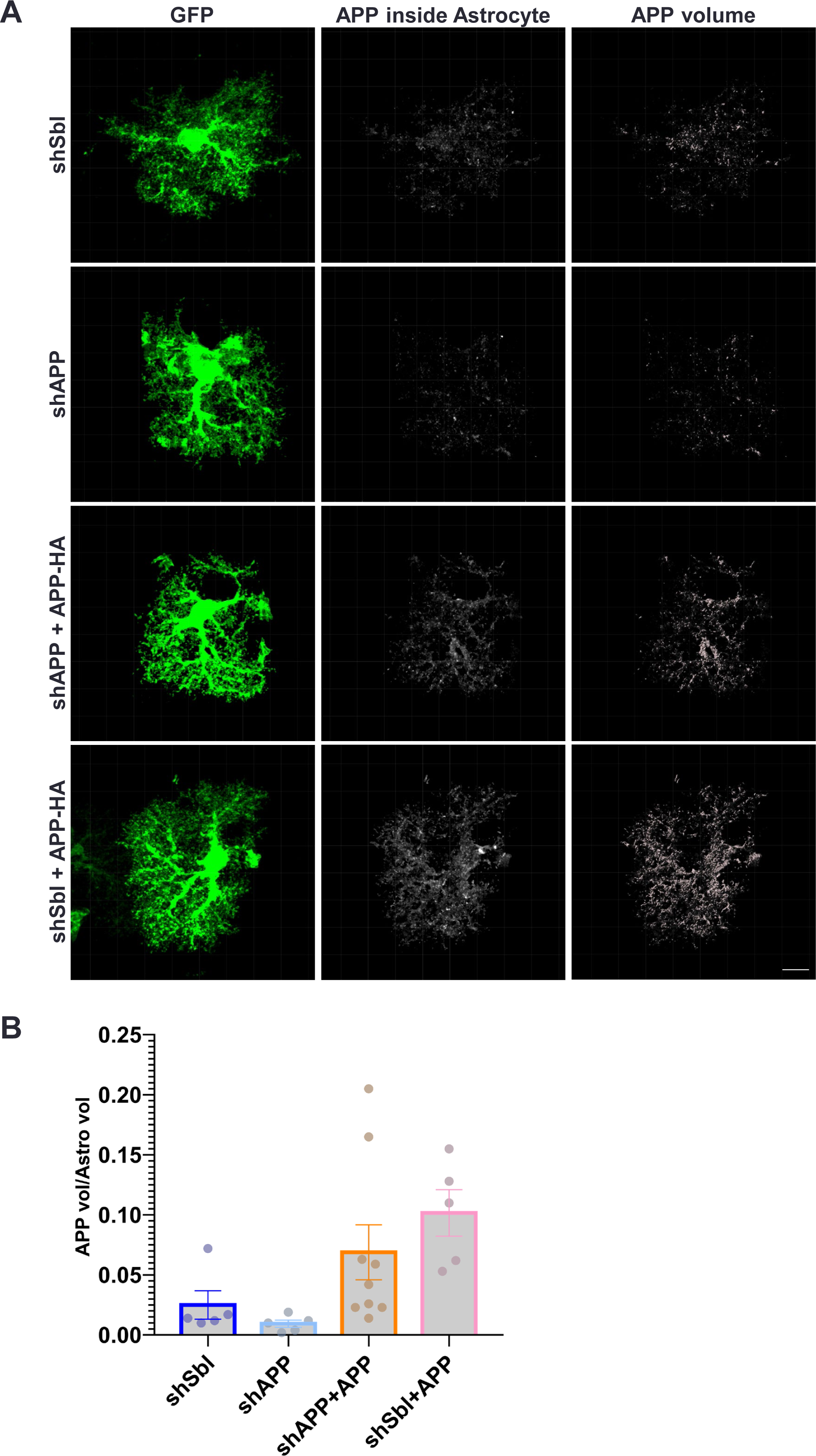
Expression of APP in cortical astrocytes overexpressing APP751. **A)** Mice were injected with AAV-pGFAP-EGFP-shSbl(miR30) or AAV-pGFAP-EGFP-shAPP(miR30) with or without AAV-pGFAPAPP751-Ct-HA. The volume of APP signal inside EGFP positive astrocytes was reconstructed and measured on IMARIS in each condition. **B)** Quantification APP volume (APP vol) per astrocyte volume (Astro vol) in shSbl, shAPP, shAPP+APP751 and shSbl+APP751 astrocytes from one experiment.

